# Sample size requirement for achieving multisite harmonization using structural brain MRI features

**DOI:** 10.1101/2022.03.12.484084

**Authors:** Pravesh Parekh, Gaurav Vivek Bhalerao, the ADBS consortium, John P John, G Venkatasubramanian

## Abstract

When data is pooled across multiple sites, the extracted features are confounded by site effects. Harmonization methods attempt to correct these site effects while preserving the biological variability within the features. However, little is known about the sample size requirement for effectively learning the harmonization parameters and their relationship with the increasing number of sites. In this study, we performed experiments to find the minimum sample size required to achieve multisite harmonization (using neuroHarmonize) using volumetric and surface features by leveraging the concept of learning curves. Our first two experiments show that site-effects are effectively removed in a univariate and multivariate manner; however, it is essential to regress the effect of covariates from the harmonized data additionally. Our following two experiments with actual and simulated data showed that the minimum sample size required for achieving harmonization grows with the increasing average Mahalanobis distances between the sites and their reference distribution. We conclude by positing a general framework to understand the site effects using the Mahalanobis distance. Further, we provide insights on the various factors in a cross-validation design to achieve optimal inter-site harmonization.

## 1. Introduction

With the advent of standardized data sharing structures such as the Brain Imaging Data Structure (Gorgolewski et al., 2016) and the availability of open-source platforms for data sharing, such as OpenNeuro (Markiewicz et al., 2021), it is increasingly common to share different kinds of neuroimaging data. Performing analyses by pooling samples across these datasets allows for increased sample size, better representation of geographical diversity, and the potential to develop robust, generalizable models. However, using multi-site data is challenging owing to factors like different scanners/hardware, differences in acquisition protocols, conditions of data acquisition (such as subject-positioning, eyes-open vs. eyes-closed during resting-state functional magnetic resonance imaging (fMRI)), variations in terms of image quality, etc. When pooling samples across scanners, it is essential to correct for site-related variations, which can otherwise influence outcome measurements like cortical thickness and brain volumes (for example, see (Fortin et al., 2018; Lee et al., 2019; Liu et al., 2020; Medawar et al., 2021; Takao et al., 2014; Wittens et al., 2021)). These “batch effects” are well-known in fields like microarray technology, where methods have been developed to correct for these effects (see (Johnson et al., 2007) and (Leek et al., 2010) for a review).

Multiple harmonization methods have been proposed for correcting scanner-related differences in neuroimaging. A regression-based procedure to correct for site-effects is to add dummy-coded scanner/site variables (for example, (Bruin et al., 2019; Fennema-Notestine et al., 2007; Pardoe et al., 2008; Rozycki et al., 2018; Segall et al., 2009; Stonnington et al., 2008). Another popular method for correcting site-specific effects is to use ComBat (Fortin et al., 2018; Johnson et al., 2007). The ComBat harmonization method models the location and scale of the variables to be harmonized, accounting for the additive and multiplicative effects of the site on the variables and preserving the site-specific biological variability (Fortin et al., 2018, 2017). Various extensions to ComBat have been proposed, such as ComBat-GAM, which models the non-linear effect of the age (Pomponio et al., 2020), CovBat, which additionally models the covariance in the data (Chen et al., n.d.), ComBat for longitudinal data (Beer et al., 2020), etc. A newly developed method, NeuroHarmony, attempts to generalize to unseen scanners/sites, taking into account the image quality metrics (Garcia-Dias et al., 2020).

Previous work has shown that harmonization methods like ComBat can remove site-related effects. For example, (Zavaliangos-Petropulu et al., 2019) examined differences in ROI-level diffusion measures and found only one remaining ROI showing significant protocol-related differences post harmonization. In (Fortin et al., 2017), the authors showed that ComBat effectively removed site-related differences from voxel-level diffusion scalar maps and ROI-level diffusion measures. Similarly, in (Fortin et al., 2018), the authors showed that the site-related effects on cortical thickness were removed using Combat. In addition to univariate results, the authors also demonstrated the removal of site-related effects in a multivariate manner: a support vector machine (SVM) classifier was unable to predict the site after harmonization.

The sample used for “learning” harmonization parameters must adequately capture site-related effects to achieve inter-site harmonization. This becomes critical in situations where harmonization needs to be carried out in a cross-validation manner – learning of the harmonization parameters happens from the training set, and these parameters are then applied to the test set (for example, in machine learning). Therefore, a central question in such paradigms is finding the minimum sample size required to eliminate the site effects. Additionally, it is essential to assess the sample size requirement in the context of a potential multivariate relationship between the variables and the site. In this paper, we attempt to address this lacuna by leveraging the concept of learning curves to find the minimum sample size required to remove site-related effects. By iteratively increasing the sample size per site and training a machine learning classifier to predict the site, we attempt to find the sample size at which the classifier prediction reduces to chance.

## 2. Methodology

### 2.1 Datasets

For this study, we selected T1-weighted MRI scans of healthy subjects from four publicly available datasets and supplemented them with scans from our labs. We restricted our selection to datasets that had scans of at least 300 healthy subjects acquired on the same scanner. The first dataset was the Southwest University Adult Lifespan Dataset (SALD) (Wei et al., 2018), a cross-sectional sample of 494 subjects. The second dataset consisted of the scans acquired at the Guy’s Hospital and available as part of the IXI dataset (available at https://brain-development.org/ixi-dataset/) and composed of 322 subjects (henceforth referred to as “Guys” dataset). The third dataset was the Amsterdam Open MRI Collection (AOMIC) (Snoek et al., 2021a, 2021b) and consisted of 928 subjects. The fourth dataset was a pooled^1^ version of four different datasets: the Beijing Normal University (BNU) dataset 1 (Lin et al., 2015) (*n* = 57; available at: https://fcon_1000.projects.nitrc.org/indi/CoRR/html/bnu_1.html), BNU dataset 2 (Huang et al., 2016) (n = 61; available at: https://fcon_1000.projects.nitrc.org/indi/CoRR/html/bnu_2.html), BNU dataset 3 (n = 48; available at: https://fcon_1000.projects.nitrc.org/indi/CoRR/html/bnu_3.html), from the Consortium for Reliability and Reproducibility (CoRR) dataset (Zuo et al., 2014), and the “Beijing_Zang” dataset (n = 198) from the 1000 Functional Connectomes Project (Biswal et al., 2010). The CoRR dataset is test-retest reliability scans, and for the present work, we only considered the baseline scans for all subjects (we henceforth refer to this combined dataset as “BNUBeijing”). The final dataset (“NIMHANS” dataset) consisted of 372 subjects collected at the National Institute of Mental Health and Neurosciences (NIMHANS) as part of two different research projects.

### 2.2 Image Acquisition and Processing

We have summarized the critical acquisition parameters for the datasets in **Table 1**. For each image, we set the origin (i.e., (0,0,0) coordinate) to correspond to the anterior commissure (AC) using *acpcdetect* v2 (Ardekani, 2018; Ardekani et al., 1997; Ardekani and Bachman, 2009) (available at: https://www.nitrc.org/projects/art/). We then, visually examined the images and manually set the origin to the AC using the display utility in SPM12 v7771 (https://www.fil.ion.ucl.ac.uk/spm/software/spm12/) in case of *acpcdetect* failure. Then, we ran the segmentation and surface pipeline for all the images using the Computational Anatomy Toolbox (CAT) 12.7 v 1727 (http://www.neuro.uni-jena.de/cat/) with SPM12 v7771 in the background, running on MATLAB R2020a (The MathWorks, Natick, USA; https://www.mathworks.com).

**Table 1:**
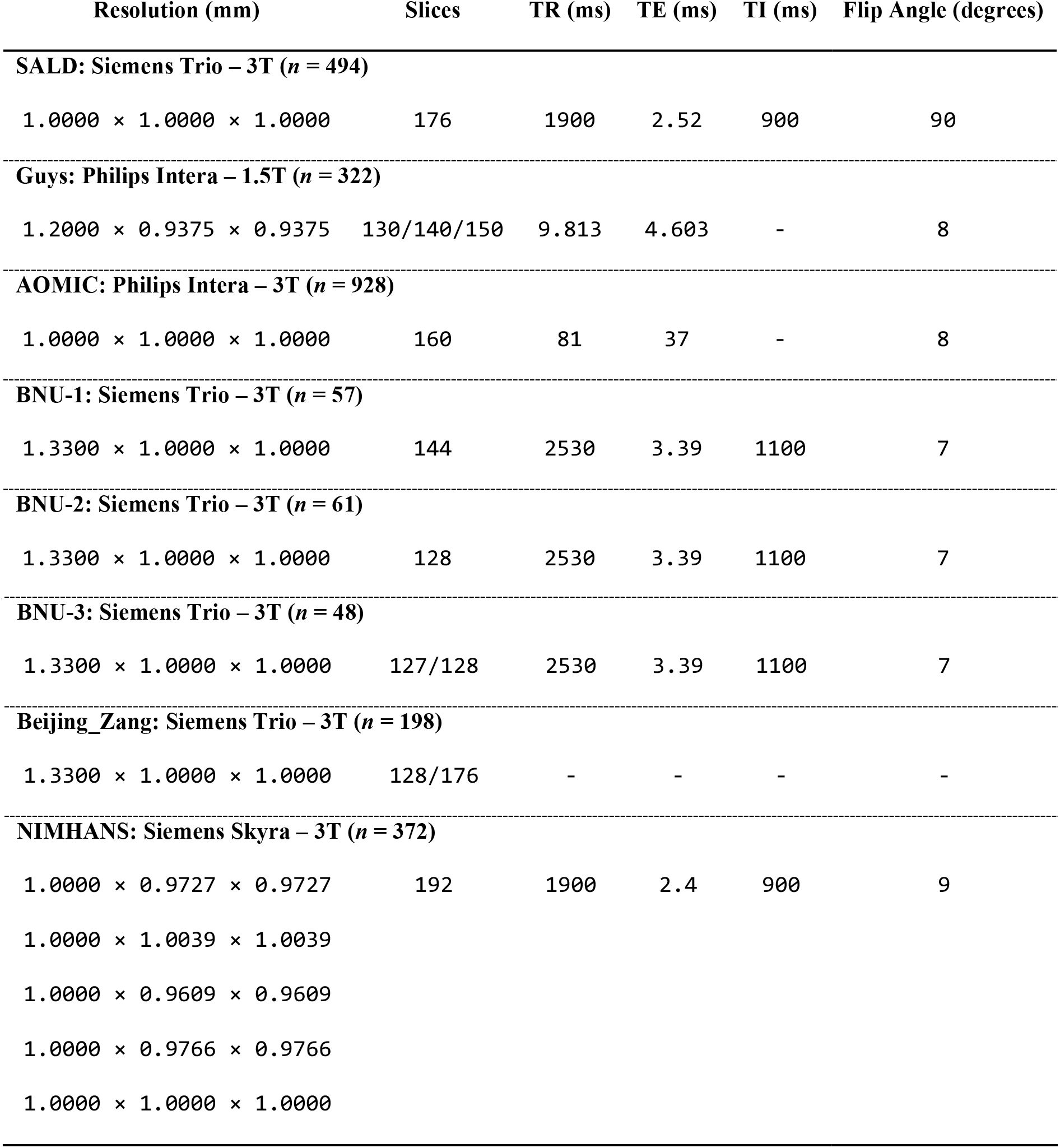
Summary of crucial acquisition parameters across the five datasets

| Resolution (mm) | Slices | TR (ms) | TE (ms) | TI (ms) | Flip Angle (degrees) |
| --- | --- | --- | --- | --- | --- |
| **SALD: Siemens Trio – 3T ($n = 494$)** |  |  |  |  |  |
| 1.0000 × 1.0000 × 1.0000 | 176 | 1900 | 2.52 | 900 | 90 |
| **Guys: Philips Intera – 1.5T ($n = 322$)** |  |  |  |  |  |
| 1.2000 × 0.9375 × 0.9375 | 130/140/150 | 9.813 | 4.603 | - | 8 |
| **AOMIC: Philips Intera – 3T ($n = 928$)** |  |  |  |  |  |
| 1.0000 × 1.0000 × 1.0000 | 160 | 81 | 37 | - | 8 |
| **BNU-1: Siemens Trio – 3T ($n = 57$)** |  |  |  |  |  |
| 1.3300 × 1.0000 × 1.0000 | 144 | 2530 | 3.39 | 1100 | 7 |
| **BNU-2: Siemens Trio – 3T ($n = 61$)** |  |  |  |  |  |
| 1.3300 × 1.0000 × 1.0000 | 128 | 2530 | 3.39 | 1100 | 7 |
| **BNU-3: Siemens Trio – 3T (*n* = 48)** |  |  |  |  |  |
| 1.3300 × 1.0000 × 1.0000 | 127/128 | 2530 | 3.39 | 1100 | 7 |
| **Beijing_Zang: Siemens Trio – 3T (*n* = 198)** |  |  |  |  |  |
| 1.3300 × 1.0000 × 1.0000 | 128/176 | - | - | - | - |
| **NIMHANS: Siemens Skyra – 3T (*n* = 372)** |  |  |  |  |  |
| 1.0000 × 0.9727 × 0.9727 | 192 | 1900 | 2.4 | 900 | 9 |
| 1.0000 × 1.0039 × 1.0039 |  |  |  |  |  |
| 1.0000 × 0.9609 × 0.9609 |  |  |  |  |  |
| 1.0000 × 0.9766 × 0.9766 |  |  |  |  |  |
| 1.0000 × 1.0000 × 1.0000 |  |  |  |  |  |

### 2.3 Quality Check

First, we rejected the data of any subject with an age of less than 18 years. Next, we flagged the data of any subject whose CAT12 report had a noise rating of “D” or below. In the next step, we eliminated the scans of any subject where the CAT12 quantified white matter hyperintensity exceeded a volume of 10 cm^3^ (while this criterion is arbitrary, it helped eliminate any scans with a potential of underlying white matter abnormalities). Additionally, to identify scans with improper segmentation, we calculated the Dice coefficient of the (binarized) modulated normalized gray matter image with the (binarized) template gray matter image (from CAT12) and flagged the images with a Dice coefficient less than 0.9. In addition, the NIMHANS dataset has undergone a thorough visual quality check as part of ongoing research work at our labs.

### 2.4 Selected Sample

After the quality check, we had 489 subjects in the SALD dataset, 302 in the Guys dataset, 928 in the AOMIC dataset, 341 in the BNUBeijing dataset, and 318 in the NIMHANS dataset. Since the lowest number was 302 subjects in the Guys dataset, we decided to randomly subset 300 subjects from each dataset for further experiments. Further, we rounded the age to the nearest integer. The socio-demographic details of these 1500 subjects are summarized in **Table 2**.

**Table 2:** Summary of socio-demographics details for each dataset after quality check

| Dataset | # Males | # Females | Age (Males):<br>Mean ± SD,<br>Min – Max | Age (Females):<br>Mean ± SD,<br>Min – Max | Age (Overall):<br>Mean ± SD,<br>Min – Max |
| --- | --- | --- | --- | --- | --- |
| SALD | 117 | 183 | 45.62 ± 17.82,<br>20 – 80 | 45.28 ± 17.46,<br>19 – 78 | 45.41 ± 17.57,<br>19 – 80 |
| Guys | 132 | 168 | 48.33 ± 16.44,<br>20 – 86 | 51.41 ± 14.96,<br>21 – 80 | 50.05 ± 15.68,<br>20 – 86 |
| AOMIC | 144 | 156 | 23.07 ± 1.74,<br>20 – 26 | 23.03 ± 1.79,<br>20 – 26 | 23.05 ± 1.77,<br>20 – 26 |
| BNUBeijing | 131 | 169 | 21.79 ± 1.86,<br>18 – 27 | 21.53 ± 1.89,<br>18 – 29 | 21.64 ± 1.88,<br>18 – 29 |
| NIMHANS | 188 | 112 | 27.20 ± 5.10,<br>18 – 49 | 27.28 ± 6.80,<br>18 – 50 | 27.23 ± 5.78,<br>18 – 50 |

### 2.5 Features

We extracted regional gray matter volumes using the Hammers atlas (CAT12 version; (Faillenot et al., 2017; Gousias et al., 2008; Hammers et al., 2003) as the primary features. This version of the Hammers atlas consists of 68 regions of interest (ROIs); from these, we excluded the bilateral parcels of corpus callosum, brainstem, and the ventricles, resulting in a total of 60 gray matter volumes for each subject. In addition to volumetric features, we also performed experiments using regional cortical thickness, fractal complexity, sulcal depth, and gyrification index. For these surface features, we used the Desikan-Killiany (DK40) atlas (Desikan et al., 2006) consisting of 72 parcellations; from these, we excluded the bilateral corpus callosum and the unknown parcels, resulting in a total of 68 surface estimates for each subject.

### 2.6 Harmonization

For all the experiments, we used the neuroHarmonize (Pomponio et al., 2020) (available at: https://github.com/rpomponio/neuroHarmonize/) toolbox for harmonizing the features across scanners. For each experiment, each model (see below), we preserved the effects of age, total intracranial volume (TIV), and sex (dummy coded as 1 for females).

### 2.7 Experiments

The primary motivation behind this study is to estimate the minimum sample size required to achieve inter-site harmonization, such that the multivariate mapping between features and scanners is removed. To achieve this, we first show that univariate (experiment 1) and multivariate (experiment 2) mapping exists between features and scanners; we also show that this mapping is removed after performing harmonization. Then, in experiment 3, we use the concept of learning curves to estimate the minimum sample size required to remove the (multivariate) site effects. Finally, in experiment 4, we extended the learning curve experiment using simulated data. This was done to examine the sample size requirement under a wide-range of Mahalanobis distances (see below) and number of sites. Critically, we note that for experiments 2-4, we used the same seeds to ensure the comparability of the results.

#### 2.7.1 Experiment 1: univariate differences

Following (Garcia-Dias et al., 2020), we performed two-sample Kolmogorov-Smirnov (KS) tests between pairs of scanners for each ROI’s brain volume and surface features to test if the distribution of the features were significantly different before and after harmonization. However, it should be noted that age, TIV, and sex will confound ROI features. Therefore, for each pair of scanners, for each ROI, we performed a linear regression to correct for these confounding variables. Then, the residuals from the linear regression were used for the pairwise two-sample KS test. Similar to the earlier work (Garcia-Dias et al., 2020), we did not perform any correction for multiple comparisons and examined our results at α = 0.05.

#### 2.7.2 Experiment 2: multivariate differences

We fit a linear SVM model to predict the scanner using volumetric features. We repeated this before and after harmonization using 10-fold cross-validation considering a pair of scanners at a time (10 combinations), three scanners at a time (10 combinations), four scanners at a time (five combinations), and all five scanners at the same time. We used the one vs. one coding method in MATLAB for multiclass classification. We performed the following operations within a cross-validation framework: regression of age, TIV, and sex (independently and combinations thereof; eight combinations), standardization of features, and training of linear SVM (using the default hyperparameter *C* = 1). In each of these steps, the parameters were learned from the training data of each fold and then applied to the test data. In the case of harmonization, the training data was harmonized, and harmonization parameters were applied to the test data before the regression step for each cross-validation fold. This was done to ensure that the regression coefficients were not confounded by site effects; a similar approach has been followed in (Pomponio et al., 2020). We repeated the 10-fold cross-validation 50 times and performed an additional 50 repeats of permutation testing (i.e., 100 repetitions for permutation testing). For permutation testing, we permuted the class labels of the entire data and calculated the *p* values as described in (Ojala and Garriga, 2010). This procedure is illustrated in **Figure 1**.

**Figure 1:**
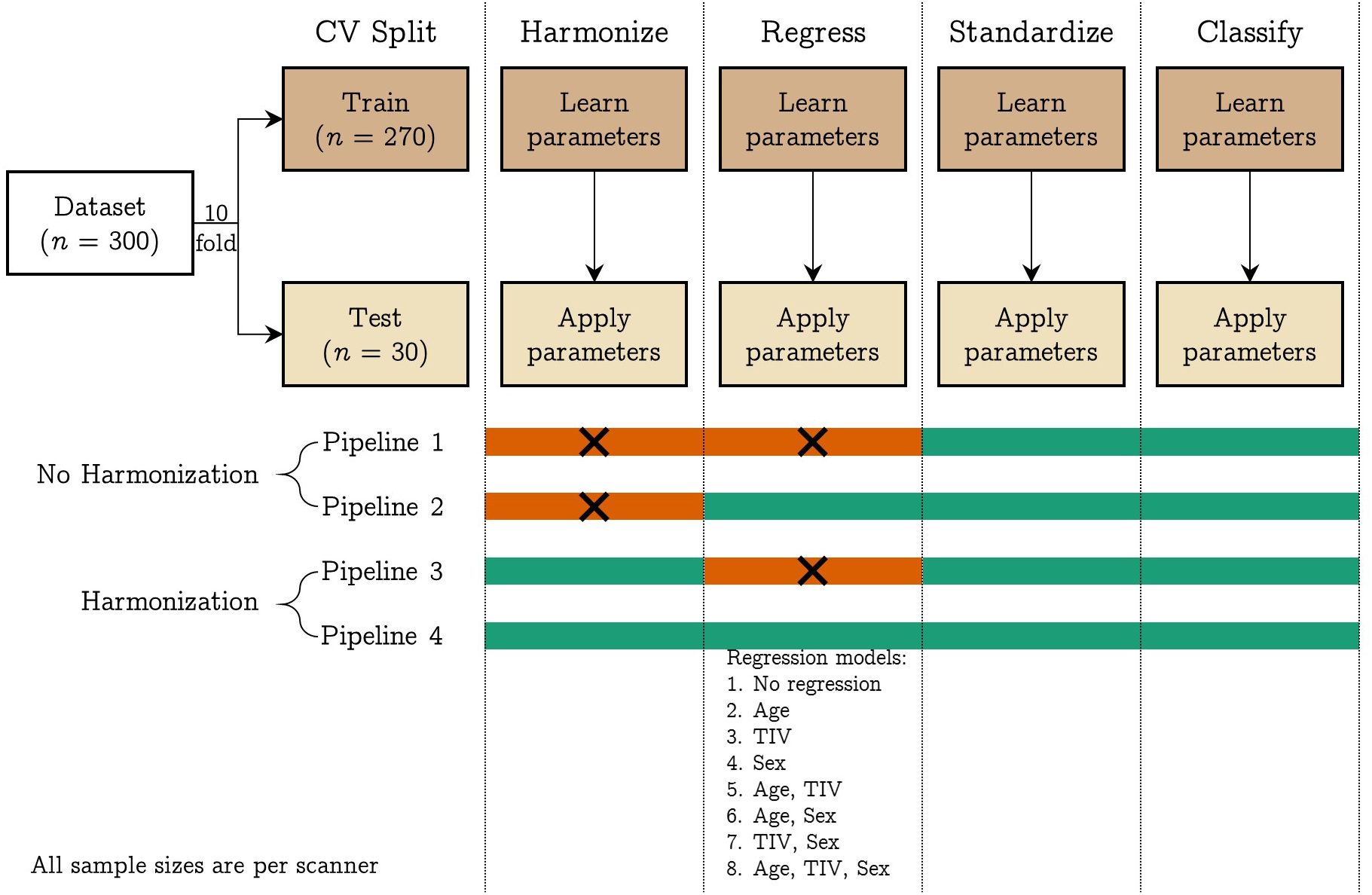
Pipelines implemented in experiment 2: we trained a linear SVM classifier to predict the scanner from raw and harmonized structural features; additionally, we explored eight different regression models where we regressed the effect of different confounding variables from the structural features; the four pipelines have four different modules: harmonization, regression, standardization, and classification; the steps indicated with orange color were not performed in that pipeline. The 10-fold cross-validation was repeated 50 times and an additional 50 repeats of permutation testing (i.e., 100 repeats of permutation) were performed to assess whether the classification performance was above chance level. [color version of this figure is available online]

#### 2.7.3 Quantifying the multivariate site-effect

In order to quantify the multivariate site-effect prevalent in our dataset, we used the Mahalanobis distance (MD) (Mahalanobis, 1936), which is a multivariate extension of Cohen’s *d* (Cohen, 1988) and can be used for calculating the standardized mean differences between groups (Del Giudice, 2009). We first calculated a reference distribution using the overall mean and pooled covariance matrix from the individual distributions. Then, we calculated the MD from each individual distribution to the reference distribution as:

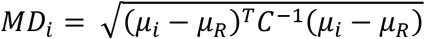

where *μ_i_* indicates the means of all variables in that individual distribution, *μ_R_* indicates the means of all variables in the reference distribution, *C* indicates the overall pooled variance-covariance matrix, and *MD_i_* indicates the calculated Mahalanobis distance of that individual distribution from the reference. We then average the individual *D_i_* s to get an overall measure of the site-effect. We calculated two versions of MD - one without regressing any covariates and one after regressing the effect of TIV, age, and sex. When performing the regression, we regressed the effect of the covariates on the entire data (i.e., linearly regressing the effect of covariates across all sites in a single model) before calculating the MD.

#### 2.7.4 Experiment 3: learning curves

In order to estimate the minimum sample size required to remove the site-effects, we leveraged the concept of learning curves. Stated simply, a learning curve can be used to assess the performance of a learner (SVM, in our case) as a function of increasing sample size. Within a cross-validation framework, when small sample sizes are used for learning harmonization parameters, we do not expect multivariate site-effects to be effectively removed from the test set. This can be tested by dividing the datasets into three parts: the first part used for learning harmonization parameters (and applying to the other parts), the second part used for learning the multivariate mapping between features and scanners, and the third part used for evaluating the classifier. If the site-effects are removed by harmonization, then a classifier will not be able to learn the mapping between features and scanners, and therefore the test accuracy of this classifier will be as good as chance level (which can be assessed by permutation testing).

The experimental design is illustrated in **Figure 2**. For this experiment, we first performed a 10-fold split on the dataset (*n* = 1500, 300 samples per scanner), resulting in 270 samples and 30 samples (“SVM test”) per scanner; the 270 samples per scanner were then split into 200 (“NH learn”) and 70 samples (“SVM train”) per scanner. The *NH learn* sample was used for learning harmonization parameters, the *SVM train* sample was used for training SVM, and the *SVM test* sample was used for testing the SVM performance. For the *NH learn* sample, we iteratively increased the sample size from 10 samples per scanner to 200 samples per scanner in increments of 10 samples (i.e., within each fold, 20 different sample sizes were used to learn harmonization parameters). For each sample size, we learnt the harmonization parameters and then applied to *SVM train* and *SVM test* samples. The harmonized *SVM train* sample was then used for training a linear SVM classifier to predict the scanner and the classifier performance assessed on harmonized *SVM test* sample. The entire process was repeated 50 times and an additional 50 repeats were performed for permutation testing (i.e., a total of 100 repeats for permutation testing). For this experiment, we learnt the regression coefficients (for the effect of age, TIV, and sex) from the *SVM train* samples and applied the coefficients to *SVM test* samples. Similarly, the standardization parameters were learnt from the *SVM train* samples and applied to the *SVM test* samples. Similar to experiment 2, we performed experiment 3 by taking a pair of scanners at a time (10 combinations), three scanners at a time (10 combinations), four scanners at a time (five combinations), and all five scanners at the same time.

**Figure 2:**
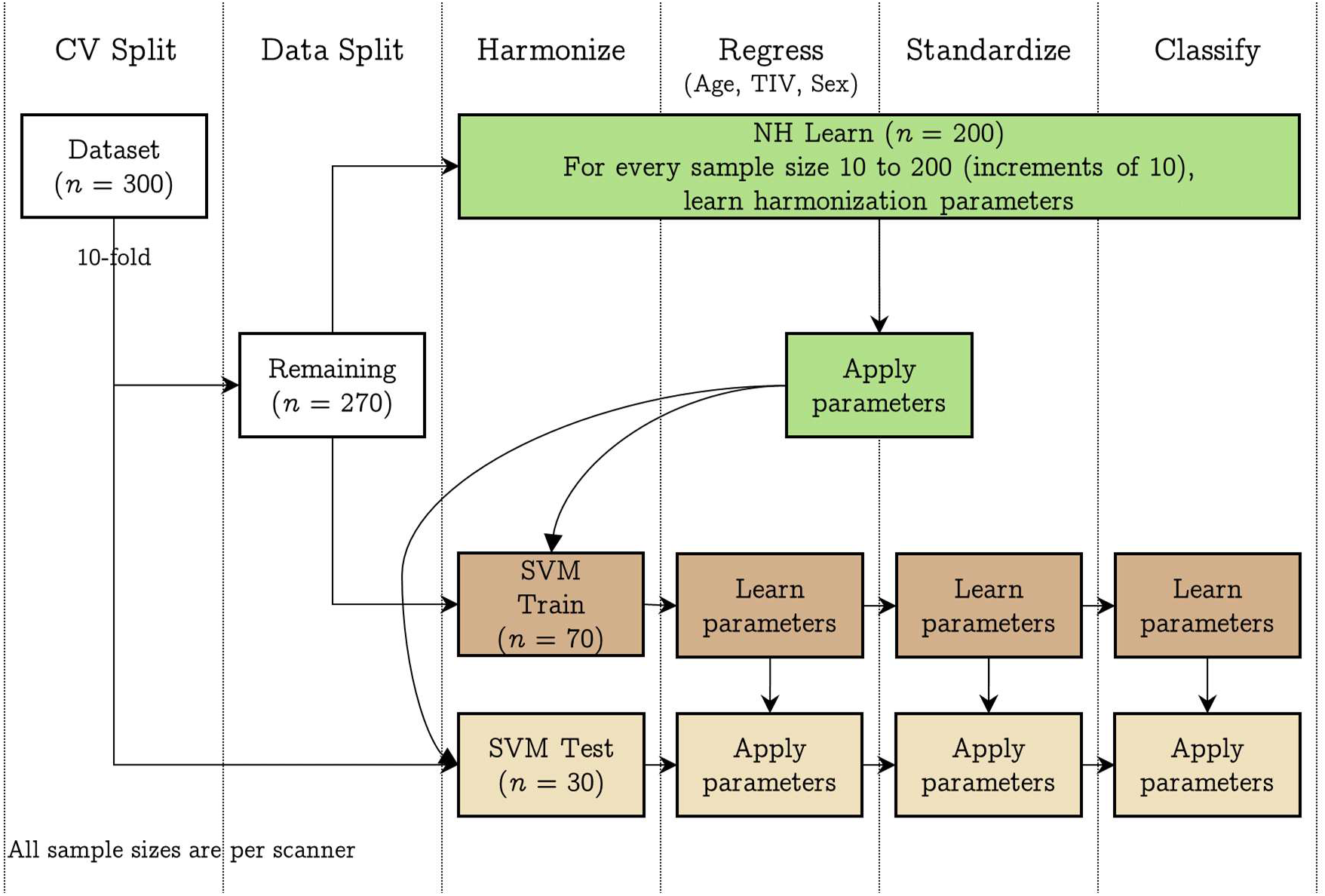
Pipeline implemented in experiment 3: we trained a linear SVM classifier to predict the scanner after using different samples sizes to achieve harmonization of structural features; first, we performed a 10-fold split on the data resulting in 30 samples per scanner (SVM Test) and 270 samples per scanner. The 270 samples were next split into 200 samples per scanner (NH learn) and 70 samples per scanner (SVM Train). For every sample size 10 to 200, at increments of 10, we learnt the harmonization parameters using NH learn and applied it to SVM Train and SVM Test samples. Then, after regressing the effect of age, TIV, and sex, we standardized the SVM Train data (and applied the regression and standardization parameters to SVM Test) and trained a linear SVM classifier to predict the scanner. Model performance was assessed on SVM Test dataset. The 10-fold cross-validation was repeated 50 times and an additional 50 repeats of permutation testing (i.e., 100 repeats of permutation) were performed to assess whether the classification performance was above chance level. [color version of this figure is available online]

#### 2.7.5 Experiment 4: simulation-based approach

We created 600 simulated samples per site by sampling from a multivariate normal distribution with means and covariances from the fractal dimension features from the actual data. Specifically, since we had five sites, we took their means and covariances and combined them to have 25 different combinations of means and covariances; using these, we obtained 25 simulated datasets having 600 samples in each site. This strategy allowed us to get datasets covering a wide range of MDs between pairs of sites. We then repeated the learning curve experiment (experiment 3) using these simulated datasets with the difference that the k-fold split was 20 fold, resulting in 30 samples per site for *SVM Test*, a holdout of 70 samples per site for *SVM Train*, and 500 samples for *NH Learn* (which we iterated over from 10 samples per site to 500 samples per site in increments of 10, stopping when the *SVM Test* performance was not above chance). The design of this experiment is shown in **Figure 3**.

**Figure 3:**
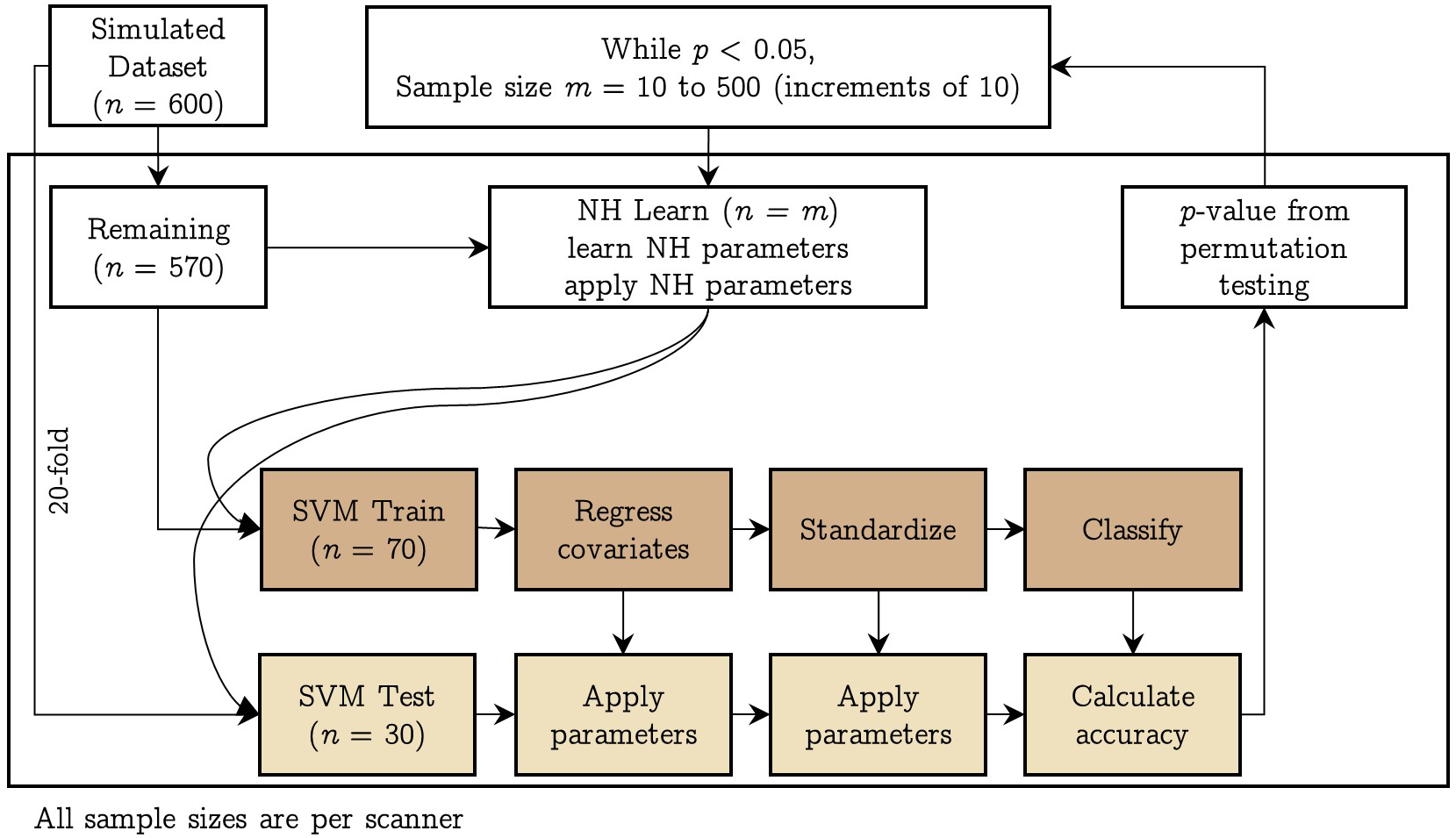
Pipeline implemented in experiment 4: using simulated data, we trained a linear SVM classifier to predict the scanner after using different samples sizes to achieve harmonization of structural features; we performed a 20-fold split on the data resulting in 30 samples per scanner (SVM Test) and 570 remaining samples per scanner. The 570 samples were next split into 500 samples per scanner (NH learn) and 70 samples per scanner (SVM Train). For every sample size 10 to 500, at increments of 10, we learnt the harmonization parameters using NH learn and applied it to SVM Train and SVM Test samples. Then, after regressing the effect of age, TIV, and sex, we standardized the SVM Train data (and applied the regression and standardization parameters to SVM Test) and trained a linear SVM classifier to predict the scanner. Model performance was assessed on SVM Test dataset. The 20-fold cross-validation was repeated 50 times and an additional 50 repeats of permutation testing (i.e., 100 repeats of permutation) were performed to assess whether the classification performance was above chance level. The whole process was repeated for every sample size in NH learn till the classification performance was above chance level. [color version of this figure is available online]

Having obtained the simulated data for 25 sites, we took combinations of two, three, and four sites taken at a time and ran experiment 3 with the data splits as mentioned above. In this simulation-based experiment, we did not attempt to simulate the covariates, in view of the complex multivariate relationship within the covariates and between the features and the covariates. Given the large number of possible combinations and the computational costs involved, we only ran the experiment by randomly^2^ selecting 100 two-, three-, and four-site combinations.

### 2.8 Data/code availability statement

The neuroimaging data used for all sites (except NIMHANS) is publicly available. The code for the experiments is available at https://github.com/parekhpravesh/HarmonizationPaper. The demographics and feature information for the NIMHANS dataset can be requested from the corresponding authors on presentation of a reasonable data analysis request. The other datasets used in this study are publicly available.

## 3. Results

### 3.1 Experiment 1: univariate difference

For every ROI, we compared the distribution of volumes and surface features between pairs of scanners before and after harmonization using a two-sample KS test at α = 0.05 (after regressing the linear effect of age, TIV, and sex). Before harmonization, several of the gray matter volumes across ROIs were statistically different between pairs of scanners; after harmonization, most of these were not significantly different, except for the right pallidum (BNUBeijing vs. NIMHANS, *p* = 0.0491) and right thalamus (SWU vs. NIMHANS, *p* = 0.0243) (see **Figure 4**). For cortical thickness, most ROIs were statistically different before harmonization; after harmonization, only left pericalcarine region was statistically significant (Guys vs. NIMHANS, *p* = 0.0113) (see Figure S1). For fractal dimension, several ROIs were significantly different before harmonization; after harmonization, the fractal dimension of only the right frontal pole was statistically significant (AOMIC vs. NIMHANS, *p* = 0.0391) (see Figure S2). For sulcal depth, several ROIs were significantly different before harmonization but after harmonization, none of the ROIs showed a statistically significant difference (see Figure S3). For gyrification index, several ROIs were significantly different before harmonization; after harmonization, the left pars orbitalis showed a statistically significant difference (AOMIC vs. BNUBeijing, *p* = 0.0309 and AOMIC vs. SWU, *p* = 0.0189) (see Figure S4). Additionally, we note that AOMIC vs. Guys and Guys vs. NIMHANS did not have many significantly different ROIs (less than 10) before harmonization for fractal dimension, sulcal depth, and gyrification index. Similarly, BNUBeijing vs. SWU only had four significantly different ROIs (fractal dimension). The overall number of significantly different ROIs before and after harmonization across feature categories is summarized in **Table 3**.

**Figure 4:**
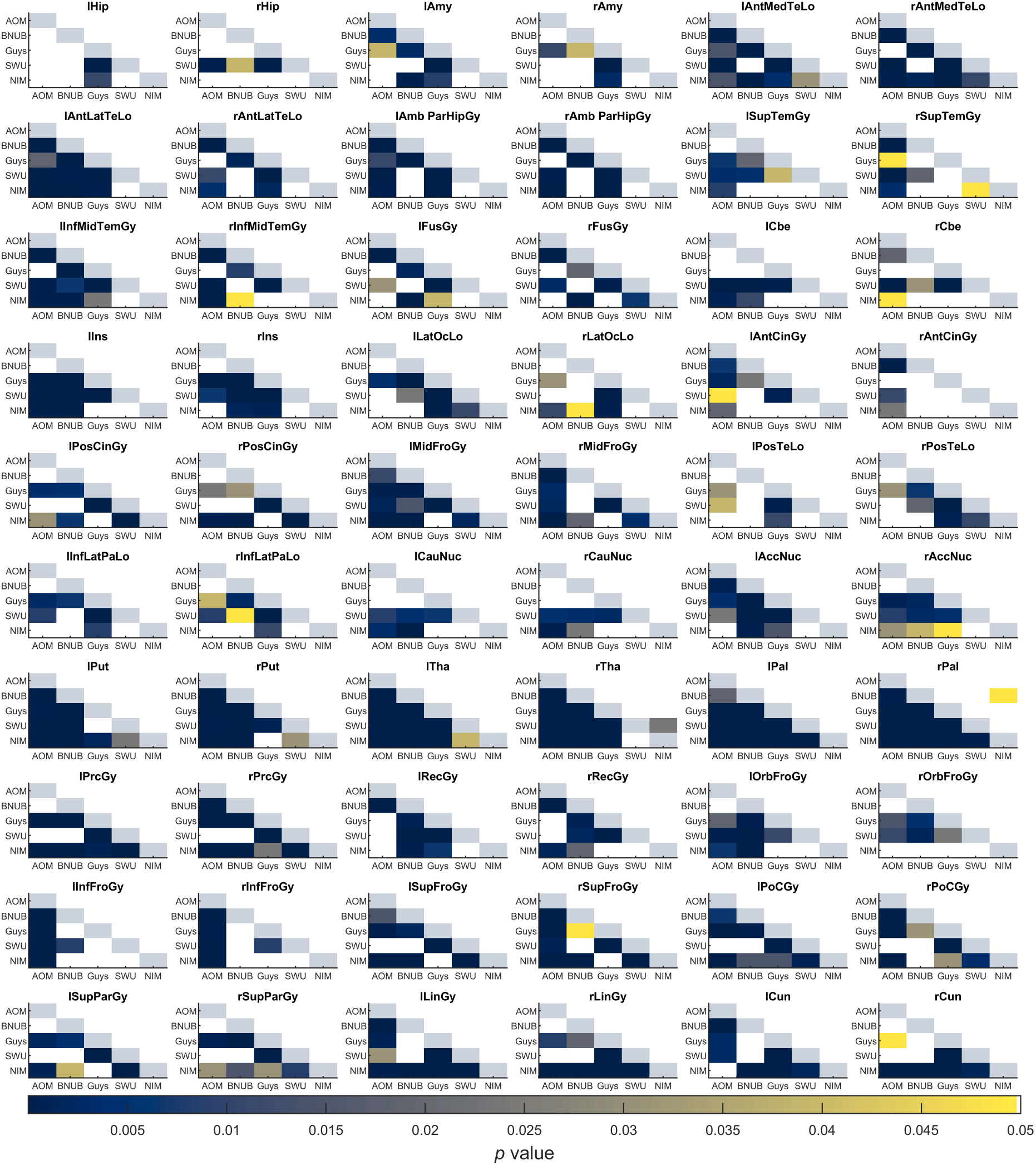
Summary of *p*-values from two sample Kolmogorov-Smirnov (KS) test between pairs of scanners for gray matter volumes. Each sub-plot indicates the *p*-values before (lower triangle) and after harmonization (upper triangle) between all pairs of scanners; the diagonal elements are shaded in a constant color to help distinguish lower and upper triangles. Each cell is color coded based on their *p*-value and only values smaller than 0.05 are shown. See Table S1 for the full names of the ROIs. Note that AOMIC dataset has been abbreviated to “AOM”, BNUBeijing dataset has been abbreviated to “BNUB”, and “NIMHANS” dataset has been abbreviated to “NIM”. [color version of this figure is available online]

**Table 3:** Number of statistically significant ROIs before and after harmonization using a two sample Kolmogorov-Smirnov (KS) test between pairs of scanners for different feature categories at α = 0.05

| Scanner | Grey matter volumes | Cortical thickness | Fractal dimension | Sulcal depth | Gyrification index |
| --- | --- | --- | --- | --- | --- |

|  | # BH | # AH | # BH | # AH | # BH | # AH | # BH | # AH | # BH | # AH |
| --- | --- | --- | --- | --- | --- | --- | --- | --- | --- | --- |
| AOMIC – BNUBeijing | 35 | 0 | 53 | 0 | 44 | 0 | 59 | 0 | 36 | 1 |
| AOMIC – Guys | 44 | 0 | 51 | 0 | 2 | 0 | 9 | 0 | 6 | 0 |
| AOMIC – SWU | 41 | 0 | 32 | 0 | 33 | 0 | 45 | 0 | 30 | 1 |
| AOMIC – NIMHANS | 44 | 0 | 29 | 0 | 23 | 1 | 31 | 0 | 13 | 0 |
| BNUBeijing – Guys | 44 | 0 | 38 | 0 | 19 | 0 | 40 | 0 | 15 | 0 |
| BNUBeijing – SWU | 28 | 0 | 35 | 0 | 4 | 0 | 18 | 0 | 13 | 0 |
| BNUBeijing – NIMHANS | 40 | 1 | 51 | 0 | 21 | 0 | 43 | 0 | 31 | 0 |
| Guys – SWU | 56 | 0 | 55 | 0 | 35 | 0 | 40 | 0 | 36 | 0 |
| Guys – NIMHANS | 35 | 0 | 47 | 1 | 8 | 0 | 8 | 0 | 7 | 0 |
| SWU – NIMHANS | 27 | 1 | 41 | 0 | 24 | 0 | 27 | 0 | 27 | 0 |

**BH:** before harmonization; **AH:** after harmonization

### 3.2 Experiment 2: multivariate differences

For volumetric features, for all site combinations, before harmonization, the SVM models were always able to predict the sites above chance level, irrespective of which covariates were regressed. When harmonization was performed (within cross-validation), for all site combinations where no regression of covariates was performed, or when TIV alone was regressed, or when TIV and sex were regressed, the SVM model predictions were above chance level. When only sex was regressed, only AOMIC vs. BNUBeijing SVM model predictions were not statistically above chance level. When age alone, or TIV and age, or age and sex were regressed, SVM prediction for certain site combinations remained statistically significant. Only when TIV, age, and sex were regressed, the SVM predictions for all site combinations were statistically not significant. An example plot showing these accuracies before and after harmonization for all categories of covariate regression (for volumetric features) is shown in **Figure 5**. Overall, all harmonized SVM accuracies were lower than before harmonization, irrespective of which covariates were regressed. This indicates that harmonization does remove site effects; however, it is important to additionally regress the confounding variables of TIV, age, and sex after harmonization to eliminate site-effects completely. For surface features, we saw a similar trend, albeit some differences (see supplementary material); the overall trend of SVM accuracies being non-significant after regression of TIV, age, and sex was consistent across the four feature categories – cortical thickness, fractal dimensions, sulcal depth, and gyrification index.

**Figure 5:**
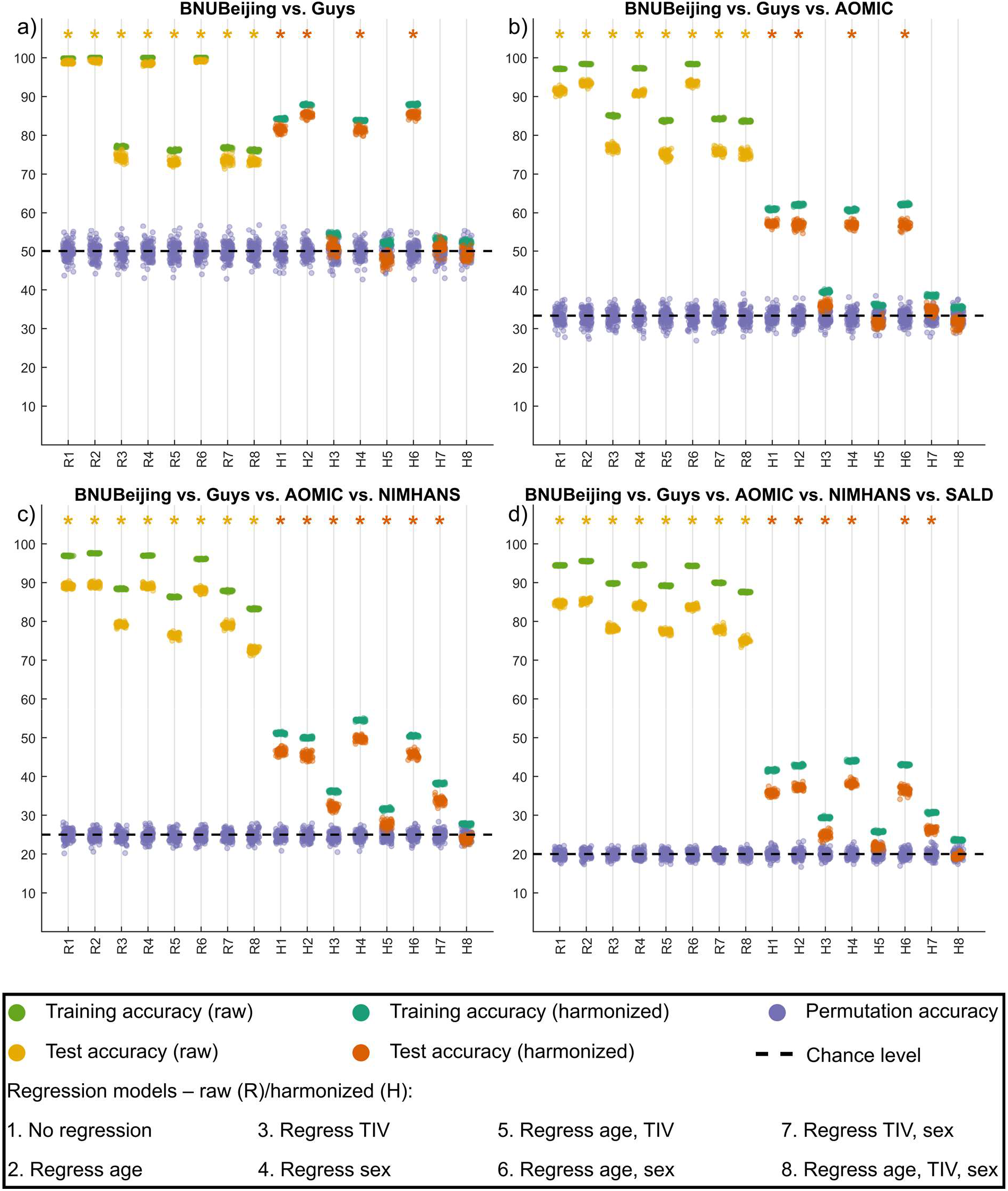
Summary of experiment 2 for representative cases for a) two sites, b) three sites, c) four sites, and d) five sites taken at a time; The *x*-axis indicates the model type – raw (R) data and harmonized (H) data with different combinations of covariates being regressed while the *y*-axis indicates the 10-fold cross-validated percentage accuracy of SVM classifier; the training and test accuracy points are across 50 repeats of 10-fold cross-validation while the permutation accuracy points are across 100 repeats of 10-fold cross-validation; the asterisk mark indicates models where the permutation testing *p*-value was less than 0.05; the theoretical chance level accuracy is indicated with a dashed black line [color version of this figure is available online]

### 3.3 Quantifying the multivariate site-effect

We used the MD to quantify the site-effect before and after regression. These results are summarized in **Figure 6**. Overall, we observed a reduction in MD after the regression of confounding variables of age, TIV, and sex. Further, for fractal dimension, sulcal depth, and gyrification index, we observed that the MD were smaller than volumetric and cortical thickness features. We observed the smallest effect sizes in SWU vs. NIMHANS (volumes), AOMIC vs. NIMHANS (cortical thickness and gyrification index), AOMIC vs. Guys (fractal dimension and sulcal depth) before regression; after regressing of the covariates, the smallest effect sizes were in AOMIC vs. Guys (for all feature categories). The largest effect sizes were seen in BNUBeijing vs. Guys (volumes), BNUBeijing vs. Guys vs. NIMHANS (cortical thickness), AOMIC vs. BNUBeijing vs. Guys vs. SWU (fractal dimension), AOMIC vs. SWU (sulcal depth), and AOMIC vs. BNUBejing vs. Guys vs. SWU vs. NIMHANS (gyrification index) before regression; after regression, the largest effect sizes were seen in AOMIC vs. BNUBeijing vs. Guys vs. SWU (volumes, cortical thickness, fractal dimension, and sulcal depth), and in AOMIC vs. BNUBeijing vs. Guys vs. SWU vs. NIMHANS (gyrification index).

**Figure 6:**
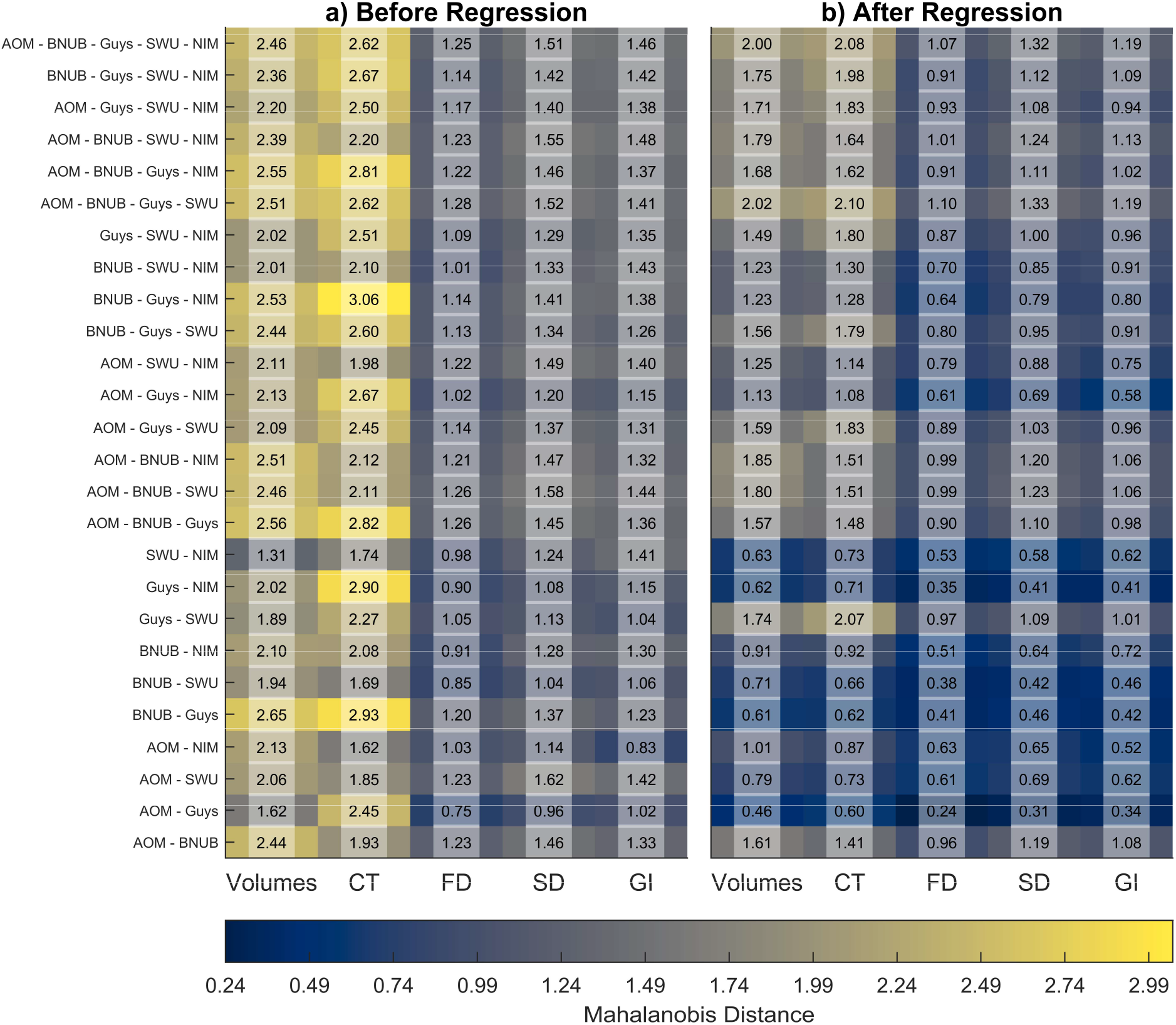
Average Mahalanobis distances between combinations of sites before and after regression of age, TIV, and sex for raw data; for any site combination, we first created a reference distribution using the overall mean and the pooled covariance; then, we calculated the distances of each site from this reference distribution and summarized it as the overall average; the x-axes indicate the different feature categories - grey matter volumes, cortical thickness (CT), fractal dimension (FD), sulcal depth (SD), and gyrification index (GI). Note that the AOMIC dataset has been abbreviated to “AOM,” BNUBeijing dataset has been abbreviated to “BNUB”, and the **NIMHANS** dataset has been abbreviated to “NIM”. [color version of this figure is available online]

### 3.4 Experiment 3: learning curves

When examining learning curves, we looked for the sample size at which the SVM classifier performance was no different than chance level (“convergence”) – we considered the sample size at which we first observed *p* >= 0.05 as the required minimum sample size to remove site-effects completely. For volumetric features, for two-site combinations, all combinations except AOMIC vs. BNUBeijing and Guys vs. SWU converged and the sample size required ranged from 120 per site (Guys vs. NIMHANS) to 180 per site (AOMIC vs. NIMHANS). When considering more than two-site combinations, we did not see convergence up to 200 samples per site (see Figures S5 and S6). The two-site results for volumetric data is summarized in **Figure 7**.

**Figure 7:**
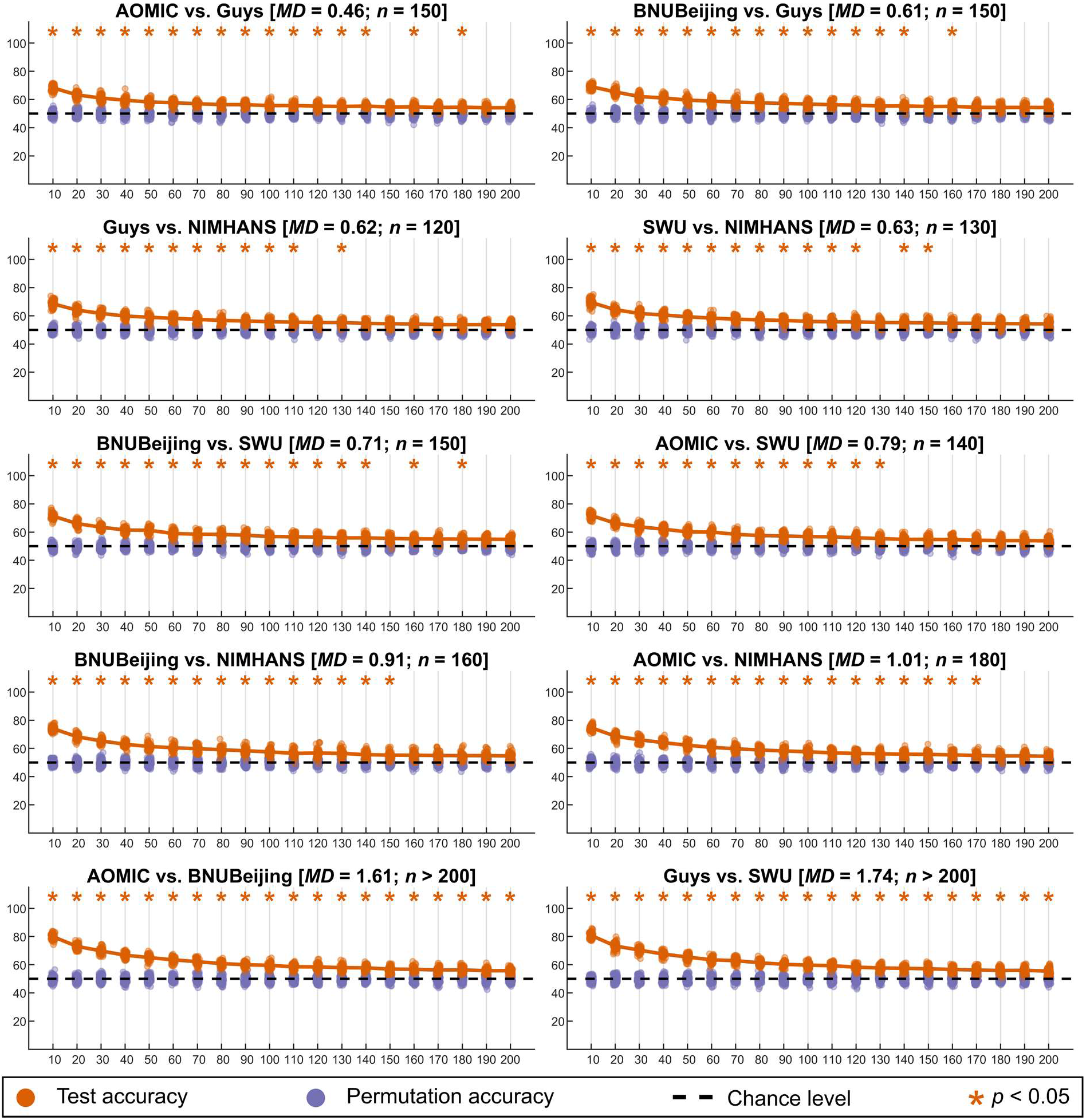
Summary of learning curves for volumetric features for two-site combinations; the orange points indicate the test accuracy of the SVM classifier (50 repeats of 10-fold cross-validation), the purple points indicate the permutation test accuracy of the SVM classifier (100 repeats of 10-fold cross-validation), while the dashed black line indicates the theoretical chance accuracy level; the x-axis indicates the sample size used for learning harmonization parameters (“*NHLearn*”) while the y-axis indicates the test accuracy in percentage. The title of each figure indicates the site-combinations, the average Mahalanobis distance (*MD*) of the two sites from the reference, and the sample size required for learning harmonization parameter (*n*) such that the SVM classifier performance was no different than chance level; the accuracies that were above chance are marked with an orange asterisk mark [color version of this figure is available online]

For cortical thickness, for two-site combinations, all combinations except BNUBeijing vs. NIMHANS and Guys vs. SWU converged (see Figure S7). The sample size required ranged from 110 per site (Guys vs. NIMHANS) to 200 per site (AOMIC vs. BNUBeijing). We did not see convergence up to 200 samples per site when considering more than two-site combinations (see Figures S8 and S9).

For fractal dimension, for two-site combinations, all combinations converged, and the sample size required ranged from 50 samples per site (AOMIC vs. Guys) to 200 samples per site (Guys vs. SWU) (see Figure S10). For three-site combinations, we observed convergence for AOMIC vs. Guys vs. NIMHANS (160 samples per site), BNUBeijing vs. SWU vs. NIMHANS (190 samples per site), and BNUBeijing vs. Guys vs. NIMHANS (200 samples per site) (see Figure S11). The other multi-site combinations did not show convergence for up to 200 samples per site (see Figure S12).

For sulcal depth, for two-site combinations, all combinations except Guys vs. SWU converged, and the required sample sizes ranged between 70 samples per site (AOMIC vs. Guys) and 160 samples per site (AOMIC vs. SWU and AOMIC vs. BNUBeijing) (see Figure S13). For three-site combinations, only AOMIC vs. Guys vs. NIMHANS converged (180 samples per site) (see Figure S14); all other multi-site combinations did not show convergence for up to 200 samples per site (see Figure S15).

For the gyrification index, for two-site combinations, all combinations except Guys vs. SWU converged, and the required sample sizes ranged between 70 samples per site (AOMIC vs. Guys) and 200 (AOMIC vs. BNUBeijing) (see Figure S16). For three-site combinations, AOMIC vs. BNUBeijing vs. Guys showed convergence at 200 samples per site (see Figure S17); all other multi-site combinations did not show convergence for up to 200 samples per site (see Figure S18).

In general, for every site-combination, we observed that the minimum sample size required for convergence increased with increasing average MDs. We observed that low MDs generally led to lower required sample size (for example, across all feature categories, the lowest MDs was ~0.24 for fractal dimension between AOMIC and Guys, which showed a convergence at a mere 50 samples per site). We calculated the correlation between the average MDs (across all feature categories) and the minimum sample sizes required per site (excluding the site combinations which did not converge) and found a strong association between these (*r* = 0.82, *p* < 0.00).

### 3.5 Experiment 4: simulation-based approach

Similar to our learning curve experiment, when examining the sample size requirement for a variety of MDs, we observed that the sample size requirement increased with increasing MDs. When comparing the sample size requirement across a number of sites for comparable MDs, we observed that the sample size requirement typically increased with an increasing number of sites. A summary of the results from the simulation-based approach is shown in **Figure 8**.

**Figure 8:**
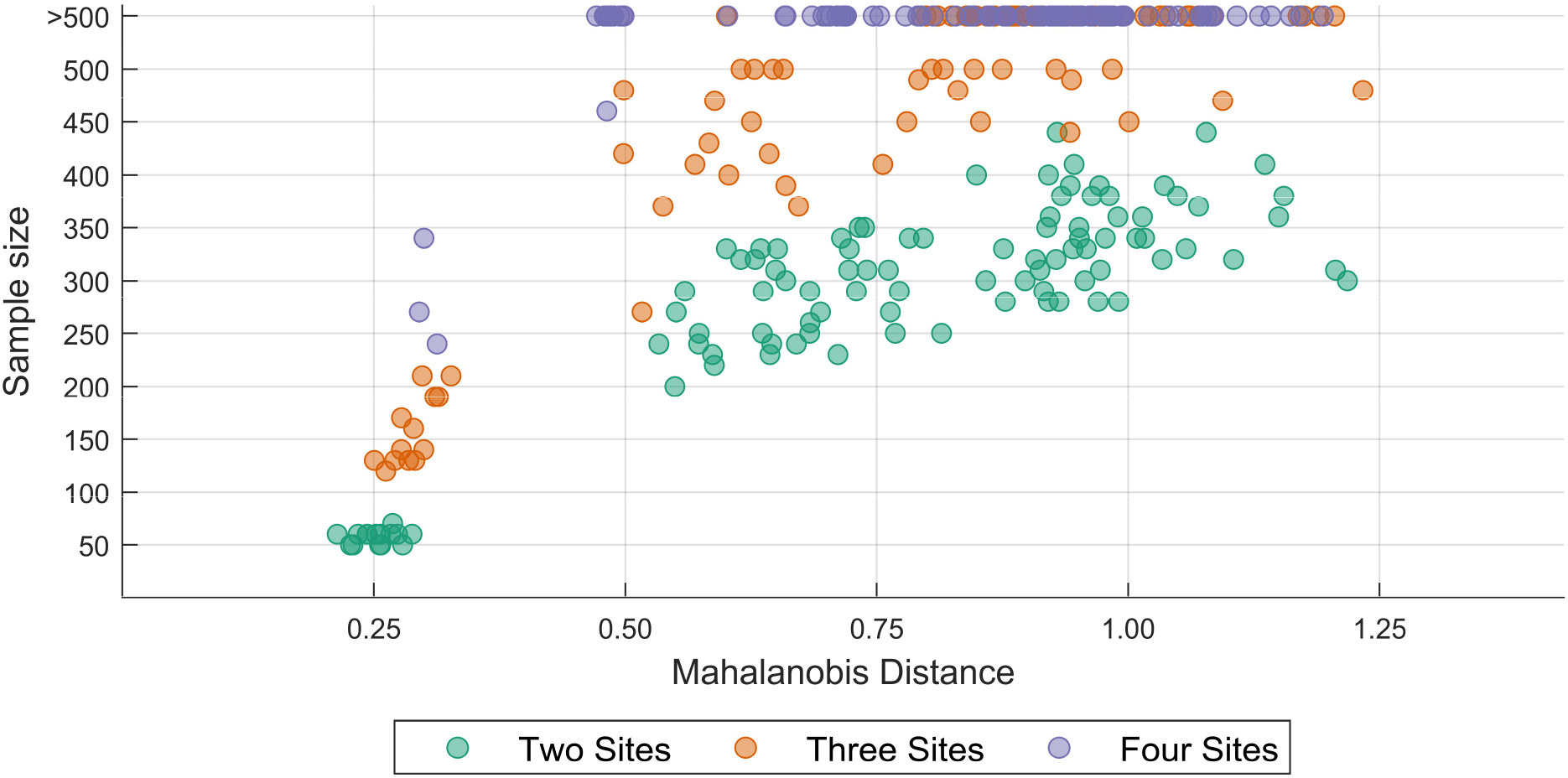
Plot of the sample size required for achieving inter-site harmonization for a range of Mahalanobis distances for two-, three-, and four-site scenarios; the features were simulated using means and covariances from fractal dimension features from real data (see text for details) [color version of this figure is available online]

## 4. Discussion

This study aimed to estimate the minimum sample size required for achieving inter-site harmonization of volumetric and surface measures. The first step in this quest was to establish whether site effects are effectively removed after harmonization. The first experiment showed that most measures across ROIs showed site-effects before harmonization; almost all the site-effects were removed (in a univariate manner) after harmonization. The few ROIs which were statistically different after harmonization are likely false positives as we did not correct for multiple comparisons. However, it is not enough to show that site-effects are corrected in a univariate manner. Therefore, we employed a machine learning approach to assess whether the site effects are eliminated in a multivariate manner. The results of this experiment showed that (in most cases) site effects are effectively removed only when performing harmonization and regressing the confounding effect of TIV, age, and sex. This reveals an interesting aspect related to the site-effect. Given that the TIV is calculated from the images, the TIV itself is confounded by site-effects. If the TIV is “preserved” during harmonization, then some residual site-effects will remain in the harmonized features. Thus, when TIV is regressed after harmonization, there is better correction of site-effects. Similarly, if other covariates are strikingly different between sites, the harmonized feature values will also show these effects. Therefore, it becomes essential to regress the effect of additional confounding factors from the harmonized data.

Next, when we examined the learning curves to find the minimum sample size needed for effective removal of site-effects (after regressing the confounding effects of TIV, age, and sex from harmonized data), we observed that the required sample size exceeded the available sample size in several cases, especially when harmonizing features from more than two sites. This is evident when examining the average MDs – as the value of MDs increases, the sample size needed to remove site-effects increases (correlation between MDs and sample size = 0.82). For example, ROI volumes and cortical thicknesses had, in general, larger values for MDs and did not show convergence for more than two sites taken at a time. In contrast, fractal dimension, on average, had smaller values for MDs and therefore showed convergence for a few of the three site combinations.

We observed a similar trend with the simulation approach, where we observed convergence at a sample size greater than 200 for larger MDs. This provides insights into why we did not see convergence (at a maximum sample size of 200) for several cases in real data. This experiment also reveals an interesting property about the sample size requirement - in general, for the same MDs, the sample size requirement increases with an increasing number of sites. However, we note that it is possible that our simulation approach overestimated the sample size requirement given that we did not simulate (and control for) the effect of covariates. In experiment 2, we showed that the site-effect reduces when covariates are controlled for - since we did not simulate and control the covariate in the simulated data, the uncorrected residual site-effects might have led to an overestimate of the sample size requirement.

### 4.1 Recommendations and future directions

#### Mahalanobis distance

In this study, we have provided a framework to examine the site-effects as the MD between each site and the reference distribution. The reference distribution is created using the overall mean and pooled covariance matrix from the individual sites. When considering more than two sites, we have computed the average of the MDs between each site and the reference distribution. However, this average MD value is just a mid-point summary statistic which will not entirely reflect the distances between the individual sites and the reference. For example, for three sites, the MD between one site and the reference may be far greater than the other two sites and the reference. Therefore, when considering the harmonization of data from more than two sites, it may additionally be useful to examine the range of the MD (i.e., the difference between the maximum MD and the minimum MD) as the sample size requirement may not only be driven by the average MD. We note that (Zhang et al., 2018) provide a data harmonization approach that allows one to specify a reference site. When using this approach, the MD can, then, be calculated with respect to the reference site rather than creating a reference distribution. In addition, future work should also examine the relationship between the minimum sample size and other distance measures. For example, distance measures which are bounded within a specific range (for example, the Jensen-Shannon divergence) could be useful in generalizing the distances across any number of sites as this might enable direct comparison of the distance measure independent of number of sites and number of features.

#### Number of sites and number of features

In our experiments, we examined the minimum sample size required for removing site effects for up to five site combinations. Further, we restricted our analyses to 60 volumetric features and 68 surface features. However, it is possible that as the number of sites and the number of features increase, the MD increases, and thereby the minimum sample size required for achieving inter-site harmonization increases. This would, of course, be dependent on the distribution of the features and the correlation among them. Future work should explore the sample size requirement by densely sampling a 3-dimensional grid of a number of sites × number of features × MDs.

#### Effect of covariates

In our second experiment, we examined the effect of preserving age, TIV, and sex as covariates and then regressed the linear effect of these covariates after harmonizing the data. These results showed that this regression step is essential to achieving complete inter-site harmonization in a multivariate sense. We have previously remarked about the relation between TIV and site effects. However, when simulating data, we have not accounted for the covariates given the multivariate relationship within the covariates and between the covariates and the features. The minimum sample size required to achieve inter-site harmonization may change depending on the number of covariates preserved during harmonization and may additionally depend on the type of the covariate (categorical/continuous), along with the MDs, the number of sites, and the number of features. Further, when harmonizing data across groups of subjects (for example, healthy subjects and patients), it might be better to have representative samples of all the groups in the harmonization training set and add the group effect as a covariate to be preserved.

#### Cross-validation

In all experiments (experiment 2-4), all the steps (including regression of covariates and standardization of data) were performed within a cross-validation framework (see **Figure 1** and **Figure 2** for experimental designs). This is an often overlooked step where confound correction is performed on the entire data before any machine learning, leading to information leakage (see, for example, (Snoek et al., 2019)). Similarly, other steps, including data harmonization and standardization, should be performed within a cross-validation framework to prevent any information leakage. Therefore, the design of the experiment (including the number of folds and the type of cross-validation) is a critical factor. It is important to ensure that the training sample is representative of the actual dataset; in addition, our results show that the number of folds is an important consideration as it will directly control the number of samples available for learning harmonization parameters.

#### Alternate methods for harmonization

As mentioned before, in addition to ComBat, other extensions to ComBat like CovBat (Chen et al., 2021) have been proposed. Similarly, recently proposed methods like NeuroHarmony (Garcia-Dias et al., 2020) need to be tested and evaluated in terms of their sample size requirement for achieving inter-site harmonization. Additionally, it would be interesting to explore the use of site-specific regression of covariates within cross-validation and mixed-effects modeling (where covariates like TIV, age, and sex are fixed-effects and site is a random-effect) as alternate methods for data harmonization.

### Conclusion

For multi-site studies that involve any form of cross-validation, it is important to carefully design the experiment such that there is enough number of samples (per site) available in the training dataset to learn the harmonization parameters adequately. Examples of such situations include machine learning classification and prediction studies and model-building studies to apply the model to new data. In this study, we have provided a framework utilizing Mahalanobis distance to quantify the site effects. Through real data and simulations, we have shown the estimated minimum sample size required to remove site effects completely. We have attempted to provide some rules of thumb for this sample size requirement under different circumstances (see **Figure 8**). However, our work indicates that further research needs to be carried out in this area while accounting for various previously enlisted factors.

## 5. Acknowledgments

This work is funded by the Department of Biotechnology, Government of India by grant number: BT/PR17316/MED/31/326/2015. The images in the “NIMHANS” dataset were acquired with funding support from Department of Science and Technology, Government of India by grant number: DST/JPJ/CSI/043 (to JPJ) and the Department of Biotechnology - Wellcome Trust India Alliance Senior Fellowship Grant (500236/Z/11/Z) (to GV). G.V. acknowledges the support of Department of Biotechnology (DBT)-Wellcome Trust India Alliance (IA/CRC/19/1/610005) and Department of Biotechnology, Government of India (BT/HRD-NBA-NWB/38/2019-20(6)). Several of the figures use the tight_subplot function (Pekka Kumpulainen (2020). tight_subplot(Nh, Nw, gap, marg_h, marg_w) (https://www.mathworks.com/matlabcentral/fileexchange/27991-tight_subplot-nh-nw-gap-marg_h-marg_w), MATLAB Central File Exchange. Retrieved January 27, 2020). The *cividis* color scheme used in **Figure 6** is from (Nuñez et al., 2018). The color schemes of several of the figures are from ColorBrewer 2.0 (http://colorbrewer2.org/) by Cynthia A. Brewer, Geography, Pennsylvania State University (accessed 25-Oct-2019).

## 6. Author contributions

**Pravesh Parekh** and **Gaurav Vivek Bhalerao**: Conceptualization, Methodology, Software, Formal analysis, Data Curation, Writing - Original Draft, Writing - Review & Editing, Visualization

**The ADBS Consortium**: Computing resources

**John P. John** and **Ganesan Venkatasubramanian:** Resources, Writing - Review & Editing, Supervision, Project administration, Funding acquisition

## 7. Ethics statement

The NIMHANS dataset was acquired as part of two research projects which received ethical clearance from the Institute Ethics Committee, prior to data collection. No additional ethical clearance was requested as the other datasets are already publicly available.

## 9. Role of funding agency

The funding agency had no role in study design, collection, analysis, and interpretation of data, writing the report, and deciding to submit the article for publication.

## Footnotes

1 We pooled these datasets as they were acquired on the same scanner; (Huang et al., 2016) mentions that the BNU series was acquired on the same scanner and Beijing_Zang was confirmed to have been acquired on the same scanner (Y.F. Zang, personal communication, December 04, 2021)

2 We randomly selected 15 MDs <= 0.5, 70 MDs between 0.5 and 1.0, and 15 MDs > 1.0

## References

1. Ardekani, B.A., 2018. A New Approach to Symmetric Registration of Longitudinal Structural MRI of the Human Brain. bioRxiv. https://doi.org/10.1101/306811

2. Ardekani, B.A., Bachman, A.H., 2009. Model-based automatic detection of the anterior and posterior commissures on MRI scans. NeuroImage 46, 677–682. https://doi.org/10.1016/j.neuroimage.2009.02.030

3. Ardekani, B.A., Kershaw, J., Braun, M., Kanuo, I., 1997. Automatic detection of the mid-sagittal plane in 3-D brain images. IEEE Transactions on Medical Imaging 16, 947–952. https://doi.org/10.1109/42.650892

4. Beer, J.C., Tustison, N.J., Cook, P.A., Davatzikos, C., Sheline, Y.I., Shinohara, R.T., Linn, K.A., 2020. Longitudinal ComBat: A method for harmonizing longitudinal multi-scanner imaging data. NeuroImage 220, 117–129. https://doi.org/10.1016/j.neuroimage.2020.117129

5. Biswal, B.B., Mennes, M., Zuo, X.-N., Gohel, S., Kelly, C., Smith, S.M., Beckmann, C.F., Adelstein, J.S., Buckner, R.L., Colcombe, S., Dogonowski, A.-M., Ernst, M., Fair, D., Hampson, M., Hoptman, M.J., Hyde, J.S., Kiviniemi, V.J., Kötter, R., Li, S.-J., Lin, C.-P., Lowe, M.J., Mackay, C., Madden, D.J., Madsen, K.H., Margulies, D.S., Mayberg, H.S., McMahon, K., Monk, C.S., Mostofsky, S.H., Nagel, B.J., Pekar, J.J., Peltier, S.J., Petersen, S.E., Riedl, V., Rombouts, S.A.R.B., Rypma, B., Schlaggar, B.L., Schmidt, S., Seidler, R.D., Siegle, G.J., Sorg, C., Teng, G.-J., Veijola, J., Villringer, A., Walter, M., Wang, L., Weng, X.-C., Whitfield-Gabrieli, S., Williamson, P., Windischberger, C., Zang, Y.-F., Zhang, H.-Y., Castellanos, F.X., Milham, M.P., 2010. Toward discovery science of human brain function. PNAS 107, 4734–4739. https://doi.org/10.1073/pnas.0911855107

6. Bruin, W., Denys, D., van Wingen, G., 2019. Diagnostic neuroimaging markers of obsessive-compulsive disorder: Initial evidence from structural and functional MRI studies. Progress in Neuro-Psychopharmacology and Biological Psychiatry, Promising neural biomarkers and predictors of treatment outcomes for psychiatric disorders: Novel neuroimaging approaches 91, 49–59. https://doi.org/10.1016/j.pnpbp.2018.08.005

7. Chen, A.A., Beer, J.C., Tustison, N.J., Cook, P.A., Shinohara, R.T., Shou, H., Initiative, T.A.D.N., 2021. Mitigating site effects in covariance for machine learning in neuroimaging data. Human Brain Mapping n/a. https://doi.org/10.1002/hbm.25688

8. Cohen, J., 1988. Statistical Power Analysis for the Behavioral Sciences, Second. ed. Routledge Academic, New York, NY. https://doi.org/10.1016/C2013-0-10517-X

9. Del Giudice, M., 2009. On the Real Magnitude of Psychological Sex Differences. Evol Psychol 7, 147470490900700220. https://doi.org/10.1177/147470490900700209

10. Desikan, R.S., Ségonne, F., Fischl, B., Quinn, B.T., Dickerson, B.C., Blacker, D., Buckner, R.L., Dale, A.M., Maguire, R.P., Hyman, B.T., Albert, M.S., Killiany, R.J., 2006. An automated labeling system for subdividing the human cerebral cortex on MRI scans into gyral based regions of interest. NeuroImage 31, 968–980. https://doi.org/10.1016/j.neuroimage.2006.01.021

11. Faillenot, I., Heckemann, R.A., Frot, M., Hammers, A., 2017. Macroanatomy and 3D probabilistic atlas of the human insula. NeuroImage 150, 88–98. https://doi.org/10.1016/j.neuroimage.2017.01.073

12. Fennema-Notestine, C., Gamst, A.C., Quinn, B.T., Pacheco, J., Jernigan, T.L., Thal, L., Buckner, R., Killiany, R., Blacker, D., Dale, A.M., Fischl, B., Dickerson, B., Gollub, R.L., 2007. Feasibility of Multi-site Clinical Structural Neuroimaging Studies of Aging Using Legacy Data. Neuroinform 5, 235–245. https://doi.org/10.1007/s12021-007-9003-9

13. Fortin, J.-P., Cullen, N., Sheline, Y.I., Taylor, W.D., Aselcioglu, I., Cook, P.A., Adams, P., Cooper, C., Fava, M., McGrath, P.J., McInnis, M., Phillips, M.L., Trivedi, M.H., Weissman, M.M., Shinohara, R.T., 2018. Harmonization of cortical thickness measurements across scanners and sites. NeuroImage 167, 104–120. https://doi.org/10.1016/j.neuroimage.2017.11.024

14. Fortin, J.-P., Parker, D., Tunç, B., Watanabe, T., Elliott, M.A., Ruparel, K., Roalf, D.R., Satterthwaite, T.D., Gur, R.C., Gur, R.E., Schultz, R.T., Verma, R., Shinohara, R.T., 2017. Harmonization of multi-site diffusion tensor imaging data. NeuroImage 161, 149–170. https://doi.org/10.1016/j.neuroimage.2017.08.047

15. Garcia-Dias, R., Scarpazza, C., Baecker, L., Vieira, S., Pinaya, W.H.L., Corvin, A., Redolfi, A., Nelson, B., Crespo-Facorro, B., McDonald, C., Tordesillas-Gutiérrez, D., Cannon, D., Mothersill, D., Hernaus, D., Morris, D., Setien-Suero, E., Donohoe, G., Frisoni, G., Tronchin, G., Sato, J., Marcelis, M., Kempton, M., van Haren, N.E.M., Gruber, O., McGorry, P., Amminger, P., McGuire, P., Gong, Q., Kahn, R.S., Ayesa-Arriola, R., van Amelsvoort, T., Ortiz-García de la Foz, V., Calhoun, V., Cahn, W., Mechelli, A., 2020. Neuroharmony: A new tool for harmonizing volumetric MRI data from unseen scanners. NeuroImage 220, 117–127. https://doi.org/10.1016/j.neuroimage.2020.117127

16. Gorgolewski, K.J., Auer, T., Calhoun, V.D., Craddock, R.C., Das, S., Duff, E.P., Flandin, G., Ghosh, S.S., Glatard, T., Halchenko, Y.O., Handwerker, D.A., Hanke, M., Keator, D., Li, X., Michael, Z., Maumet, C., Nichols, B.N., Nichols, T.E., Pellman, J., Poline, J.-B., Rokem, A., Schaefer, G., Sochat, V., Triplett, W., Turner, J.A., Varoquaux, G., Poldrack, R.A., 2016. The brain imaging data structure, a format for organizing and describing outputs of neuroimaging experiments. Scientific Data 3, 160044. https://doi.org/10.1038/sdata.2016.44

17. Gousias, I.S., Rueckert, D., Heckemann, R.A., Dyet, L.E., Boardman, J.P., Edwards, A.D., Hammers, A., 2008. Automatic segmentation of brain MRIs of 2-year-olds into 83 regions of interest. NeuroImage 40, 672–684. https://doi.org/10.1016/j.neuroimage.2007.11.034

18. Hammers, A., Allom, R., Koepp, M.J., Free, S.L., Myers, R., Lemieux, L., Mitchell, T.N., Brooks, D.J., Duncan, J.S., 2003. Three-dimensional maximum probability atlas of the human brain, with particular reference to the temporal lobe. Hum. Brain Mapp. 19, 224–247. https://doi.org/10.1002/hbm.10123

19. Huang, L., Huang, T., Zhen, Z., Liu, J., 2016. A test-retest dataset for assessing long-term reliability of brain morphology and resting-state brain activity. Sci Data 3, 160016. https://doi.org/10.1038/sdata.2016.16

20. Johnson, W.E., Li, C., Rabinovic, A., 2007. Adjusting batch effects in microarray expression data using empirical Bayes methods. Biostatistics 8, 118–127. https://doi.org/10.1093/biostatistics/kxj037

21. Lee, H., Nakamura, K., Narayanan, S., Brown, R.A., Arnold, D.L., 2019. Estimating and accounting for the effect of MRI scanner changes on longitudinal whole-brain volume change measurements. NeuroImage 184, 555–565. https://doi.org/10.1016/j.neuroimage.2018.09.062

22. Leek, J.T., Scharpf, R.B., Bravo, H.C., Simcha, D., Langmead, B., Johnson, W.E., Geman, D., Baggerly, K., Irizarry, R.A., 2010. Tackling the widespread and critical impact of batch effects in high-throughput data. Nat Rev Genet 11, 733–739. https://doi.org/10.1038/nrg2825

23. Lin, Q., Dai, Z., Xia, M., Han, Z., Huang, R., Gong, G., Liu, C., Bi, Y., He, Y., 2015. A connectivity-based test-retest dataset of multi-modal magnetic resonance imaging in young healthy adults. Sci Data 2, 150056. https://doi.org/10.1038/sdata.2015.56

24. Liu, S., Hou, B., Zhang, Y., Lin, T., Fan, X., You, H., Feng, F., 2020. Inter-scanner reproducibility of brain volumetry: influence of automated brain segmentation software. BMC Neuroscience 21, 35. https://doi.org/10.1186/s12868-020-00585-1

25. Mahalanobis, P.C., 1936. On the generalized distance in statistics. Proceedings of the National Institute of Sciences (Calcutta) 2, 49–55.

26. Markiewicz, C.J., Gorgolewski, K.J., Feingold, F., Blair, R., Halchenko, Y.O., Miller, E., Hardcastle, N., Wexler, J., Esteban, O., Goncavles, M., Jwa, A., Poldrack, R., 2021. The OpenNeuro resource for sharing of neuroscience data. eLife 10, e71774. https://doi.org/10.7554/eLife.71774

27. Medawar, E., Thieleking, R., Manuilova, I., Paerisch, M., Villringer, A., Witte, A.V., Beyer, F., 2021. Estimating the effect of a scanner upgrade on measures of grey matter structure for longitudinal designs. PLOS ONE 16, e0239021. https://doi.org/10.1371/journal.pone.0239021

28. Nuñez, J.R., Anderton, C.R., Renslow, R.S., 2018. Optimizing colormaps with consideration for color vision deficiency to enable accurate interpretation of scientific data. PLOS ONE 13, e0199239. https://doi.org/10.1371/journal.pone.0199239

29. Ojala, M., Garriga, G.C., 2010. Permutation Tests for Studying Classifier Performance. Journal of Machine Learning Research 11, 1833–1863.

30. Pardoe, H., Pell, G.S., Abbott, D.F., Berg, A.T., Jackson, G.D., 2008. Multi-site voxel-based morphometry: Methods and a feasibility demonstration with childhood absence epilepsy. NeuroImage 42, 611–616. https://doi.org/10.1016/j.neuroimage.2008.05.007

31. Pomponio, R., Erus, G., Habes, M., Doshi, J., Srinivasan, D., Mamourian, E., Bashyam, V., Nasrallah, I.M., Satterthwaite, T.D., Fan, Y., Launer, L.J., Masters, C.L., Maruff, P., Zhuo, C., Völzke, H., Johnson, S.C., Fripp, J., Koutsouleris, N., Wolf, D.H., Gur, Raquel, Gur, Ruben, Morris, J., Albert, M.S., Grabe, H.J., Resnick, S.M., Bryan, R.N., Wolk, D.A., Shinohara, R.T., Shou, H., Davatzikos, C., 2020. Harmonization of large MRI datasets for the analysis of brain imaging patterns throughout the lifespan. NeuroImage 208, 116450. https://doi.org/10.1016/j.neuroimage.2019.116450

32. Rozycki, M., Satterthwaite, T.D., Koutsouleris, N., Erus, G., Doshi, J., Wolf, D.H., Fan, Y., Gur, R.E., Gur, R.C., Meisenzahl, E.M., Zhuo, C., Yin, H., Yan, H., Yue, W., Zhang, D., Davatzikos, C., 2018. Multisite Machine Learning Analysis Provides a Robust Structural Imaging Signature of Schizophrenia Detectable Across Diverse Patient Populations and Within Individuals. Schizophr Bull 44, 1035–1044. https://doi.org/10.1093/schbul/sbx137

33. Segall, J.M., Turner, J.A., van Erp, T.G.M., White, T., Bockholt, H.J., Gollub, R.L., Ho, B.C., Magnotta, V., Jung, R.E., McCarley, R.W., Schulz, S.C., Lauriello, J., Clark, V.P., Voyvodic, J.T., Diaz, M.T., Calhoun, V.D., 2009. Voxel-based Morphometric Multisite Collaborative Study on Schizophrenia. Schizophr Bull 35, 82–95. https://doi.org/10.1093/schbul/sbn150

34. Snoek, L., Miletić, S., Scholte, H.S., 2019. How to control for confounds in decoding analyses of neuroimaging data. NeuroImage 184, 741–760. https://doi.org/10.1016/j.neuroimage.2018.09.074

35. Snoek, L., van der Miesen, M.M., Beemsterboer, T., van der Leij, A., Eigenhuis, A., Scholte, S.H., 2021a. The Amsterdam Open MRI Collection, a set of multimodal MRI datasets for individual difference analyses. Sci Data 8, 85. https://doi.org/10.1038/s41597-021-00870-6

36. Snoek, L., van der Miesen, M.M., van der Leij, A., Beemsterboer, T., Eigenhuis, A., Scholte, S.H., 2021b. AOMIC-ID1000.

37. Stonnington, C.M., Tan, G., Klöppel, S., Chu, C., Draganski, B., Jack, C.R., Chen, K., Ashburner, J., Frackowiak, R.S.J., 2008. Interpreting scan data acquired from multiple scanners: A study with Alzheimer’s disease. NeuroImage 39, 1180–1185. https://doi.org/10.1016/j.neuroimage.2007.09.066

38. Takao, H., Hayashi, N., Ohtomo, K., 2014. Effects of study design in multi-scanner voxel-based morphometry studies. NeuroImage 84, 133–140. https://doi.org/10.1016/j.neuroimage.2013.08.046

39. Wei, D., Zhuang, K., Ai, L., Chen, Q., Yang, W., Liu, W., Wang, K., Sun, J., Qiu, J., 2018. Structural and functional brain scans from the cross-sectional Southwest University adult lifespan dataset. Sci Data 5, 180134. https://doi.org/10.1038/sdata.2018.134

40. Wittens, M.M.J., Allemeersch, G.-J., Sima, D.M., Naeyaert, M., Vanderhasselt, T., Vanbinst, A.-M., Buls, N., De Brucker, Y., Raeymaekers, H., Fransen, E., Smeets, D., van Hecke, W., Nagels, G., Bjerke, M., de Mey, J., Engelborghs, S., 2021. Inter-and Intra-Scanner Variability of Automated Brain Volumetry on Three Magnetic Resonance Imaging Systems in Alzheimer’s Disease and Controls. Frontiers in Aging Neuroscience 13, 641. https://doi.org/10.3389/fnagi.2021.746982

41. Zavaliangos-Petropulu, A., Nir, T.M., Thomopoulos, S.I., Reid, R.I., Bernstein, M.A., Borowski, B., Jack Jr., C.R., Weiner, M.W., Jahanshad, N., Thompson, P.M., 2019. Diffusion MRI Indices and Their Relation to Cognitive Impairment in Brain Aging: The Updated Multi-protocol Approach in ADNI3. Frontiers in Neuroinformatics 13, 2. https://doi.org/10.3389/fninf.2019.00002

42. Zhang, Y., Jenkins, D.F., Manimaran, S., Johnson, W.E., 2018. Alternative empirical Bayes models for adjusting for batch effects in genomic studies. BMC Bioinformatics 19, 262. https://doi.org/10.1186/s12859-018-2263-6

43. Zuo, X.-N., Anderson, J.S., Bellec, P., Birn, R.M., Biswal, B.B., Blautzik, J., Breitner, J.C.S., Buckner, R.L., Calhoun, V.D., Castellanos, F.X., Chen, A., Chen, B., Chen, J., Chen, X., Colcombe, S.J., Courtney, W., Craddock, R.C., Di Martino, A., Dong, H.-M., Fu, X., Gong, Q., Gorgolewski, K.J., Han, Y., He, Ye, He, Yong, Ho, E., Holmes, A., Hou, X.-H., Huckins, J., Jiang, T., Jiang, Y., Kelley, W., Kelly, C., King, M., LaConte, S.M., Lainhart, J.E., Lei, X., Li, H.-J., Li, Kaiming, Li, Kuncheng, Lin, Q., Liu, D., Liu, J., Liu, X., Liu, Y., Lu, G., Lu, J., Luna, B., Luo, J., Lurie, D., Mao, Y., Margulies, D.S., Mayer, A.R., Meindl, T., Meyerand, M.E., Nan, W., Nielsen, J.A., O’Connor, D., Paulsen, D., Prabhakaran, V., Qi, Z., Qiu, J., Shao, C., Shehzad, Z., Tang, W., Villringer, A., Wang, H., Wang, K., Wei, D., Wei, G.-X., Weng, X.-C., Wu, X., Xu, T., Yang, N., Yang, Z., Zang, Y.-F., Zhang, L., Zhang, Q., Zhang, Zhe, Zhang, Zhiqiang, Zhao, K., Zhen, Z., Zhou, Y., Zhu, X.-T., Milham, M.P., 2014. An open science resource for establishing reliability and reproducibility in functional connectomics. Sci Data 1, 140049. https://doi.org/10.1038/sdata.2014.49

