## Supplementary Material for "Sample size requirement for achieving multisite harmonization using structural brain MRI features"

Parekh, Pravesh [a, c, d, *]

Bhalerao, Gaurav Vivek [b, c, d, *]

the ADBS consortium^

John, John P [a, c, d, #]

Venkatasubramanian, G [b, c, d, #]

* equal contribution; [#] Corresponding authors

**Keywords:** neuroimaging; harmonization; sample size; multisite; Mahalanobis distance; cross-validation

**Affiliations:**

[a] Multimodal Brain Image Analysis Laboratory, National Institute of Mental Health and Neurosciences (NIMHANS), Bangalore, India

[b] Translational Psychiatry Lab, National Institute of Mental Health and Neurosciences (NIMHANS), Bangalore, India

[c] ADBS Neuroimaging Centre, National Institute of Mental Health and Neurosciences (NIMHANS), Bangalore, India

[d] Department of Psychiatry, National Institute of Mental Health and Neurosciences (NIMHANS), Bangalore, India

^Accelerator Program for Discovery in Brain disorders using Stem cells (ADBS) Consortium:

Biju Viswanath[1], Naren P. Rao[1], Janardhanan C. Narayanaswamy[1], Palanimuthu T. Sivakumar[1], Arun Kandasamy[1], Muralidharan Kesavan[1], Urvakhsh Meherwan Mehta[1], Odity Mukherjee[2], Meera Purushottam[1], Ramakrishnan Kannan[1], Bhupesh Mehta[1], Thennarasu Kandavel[1], B. Binukumar[1], Jitender Saini[1], Deepak Jayarajan[1], A. Shyamsundar[1], Sydney Moirangthem[1], K. G. Vijay Kumar[1], Jayant Mahadevan[1], Bharath Holla[1], Jagadisha Thirthalli[1], Prabha S. Chandra[1], Bangalore N. Gangadhar[1], Pratima Murthy[1], Mitradas M. Panicker[3], Upinder S. Bhalla[3], Sumantra Chattarji[3], Vivek Benegal[1], Mathew Varghese[1], Janardhan Y. C. Reddy[1], Padinjat Raghu[3,] Mahendra Rao[3], and Sanjeev Jain

[1]National Institute of Mental Health and Neurosciences (NIMHANS); [2]Institute for Stem Cell Biology and Regenerative Medicine (InStem); [3]National Center for Biological Sciences (NCBS)

**Address for correspondence:**

Dr. John P. John
Multimodal Brain Image Analysis Laboratory
Department of Psychiatry,
National Institute of Mental Health and Neurosciences (NIMHANS),
Bangalore - 560029, India

(+91) 080 2699 5329

Dr. Ganesan Venkatasubramanian
Translational Psychiatry Lab
Department of Psychiatry,
National Institute of Mental Health and Neurosciences (NIMHANS),
Bangalore - 560029, India

(+91) 080 2699 5256

### List of ROIs: Hammers atlas

**Table S1:** Abbreviations and full names of the 60 regions of interest from the Hammers atlas; note that the abbreviation and the full names are as specified in the Computational Anatomy Toolbox (CAT)

| Abbreviation | ROI Name |
|---|---|
| lHip | Left Hippocampus |
| rHip | Right Hippocampus |
| lAmy | Left Amygdala |
| rAmy | Right Amygdala |
| lAntMedTeLo | Left Anterior Medial Temporal Lobe |
| rAntMedTeLo | Right Anterior Medial Temporal Lobe |
| lAntLatTeLo | Left Anterior Lateral Temporal Lobe |
| rAntLatTeLo | Right Anterior Lateral Temporal Lobe |
| lAmb ParHipGy | Left Ambient and Parahippocampus Gyri |
| rAmb ParHipGy | Right Ambient and Parahippocampus Gyri |
| lSupTemGy | Left Superior Temporal Gyrus |
| rSupTemGy | Right Superior Temporal Gyrus |
| lInfMidTemGy | Left Inferior Middle Temporal Gyri |
| rInfMidTemGy | Right Inferior Middle Temporal Gyri |
| lFusGy | Left Fusiform Gyrus |
| rFusGy | Right Fusiform Gyrus |
| lCbe | Left Cerebellum |
| rCbe | Right Cerebellum |
| lIns | Left Insula |
| rIns | Right Insula |
| lLatOcLo | Left Lateral Occipital Lobe |
| rLatOcLo | Right Lateral Occipital Lobe |
| lAntCinGy | Left Anterior Cinguli Gyrus |
| rAntCinGy | Right Anterior Cinguli Gyrus |
| lPosCinGy | Left Posterior Cinguli Gyrus |
| rPosCinGy | Right Posterior Cinguli Gyrus |
| lMidFroGy | Left Middle Frontal Gyrus |
| rMidFroGy | Right Middle Frontal Gyrus |
| lPosTeLo | Left Posterior Temporal Lobe |
| rPosTeLo | Right Posterior Temporal Lobe |
| lInfLatPaLo | Left Inferior Lateral Pariatal Lobe |
| rInfLatPaLo | Right Inferior Lateral Pariatal Lobe |
| lCauNuc | Left Caudate Nucleus |
| rCauNuc | Right Caudate Nucleus |
| lAccNuc | Left Accumbens Nucleus |
| rAccNuc | Right Accumbens Nucleus |
| lPut | Left Putamen |
| rPut | Right Putamen |
| lTha | Left Thalamus |
| rTha | Right Thalamus |
| lPal | Left Pallidum |
| rPal | Right Pallidum |
| lPrcGy | Left Precentral Gyrus |
| rPrcGy | Right Precentral Gyrus |
| lRecGy | Left Gyrus Rectus |

| | |
|---|---|
| rRecGy | Right Gyrus Rectus |
| lOrbFroGy | Left Orbito-Frontal Gyri |
| rOrbFroGy | Right Orbito-Frontal Gyri |
| lInfFroGy | Left Inferior Frontal Gyrus |
| rInfFroGy | Right Inferior Frontal Gyrus |
| lSupFroGy | Left Superior Frontal Gyrus |
| rSupFroGy | Right Superior Frontal Gyrus |
| lPoCGy | Left Postcentral Gyrus |
| rPoCGy | Right Postcentral Gyrus |
| lSupParGy | Left Superior Parietal Gyrus |
| rSupParGy | Right Superior Parietal Gyrus |
| lLinGy | Left Lingual Gyrus |
| rLinGy | Right Lingual Gyrus |
| lCun | Left Cuneus |
| rCun | Right Cuneus |

### Experiment 1: univariate differences – cortical thickness

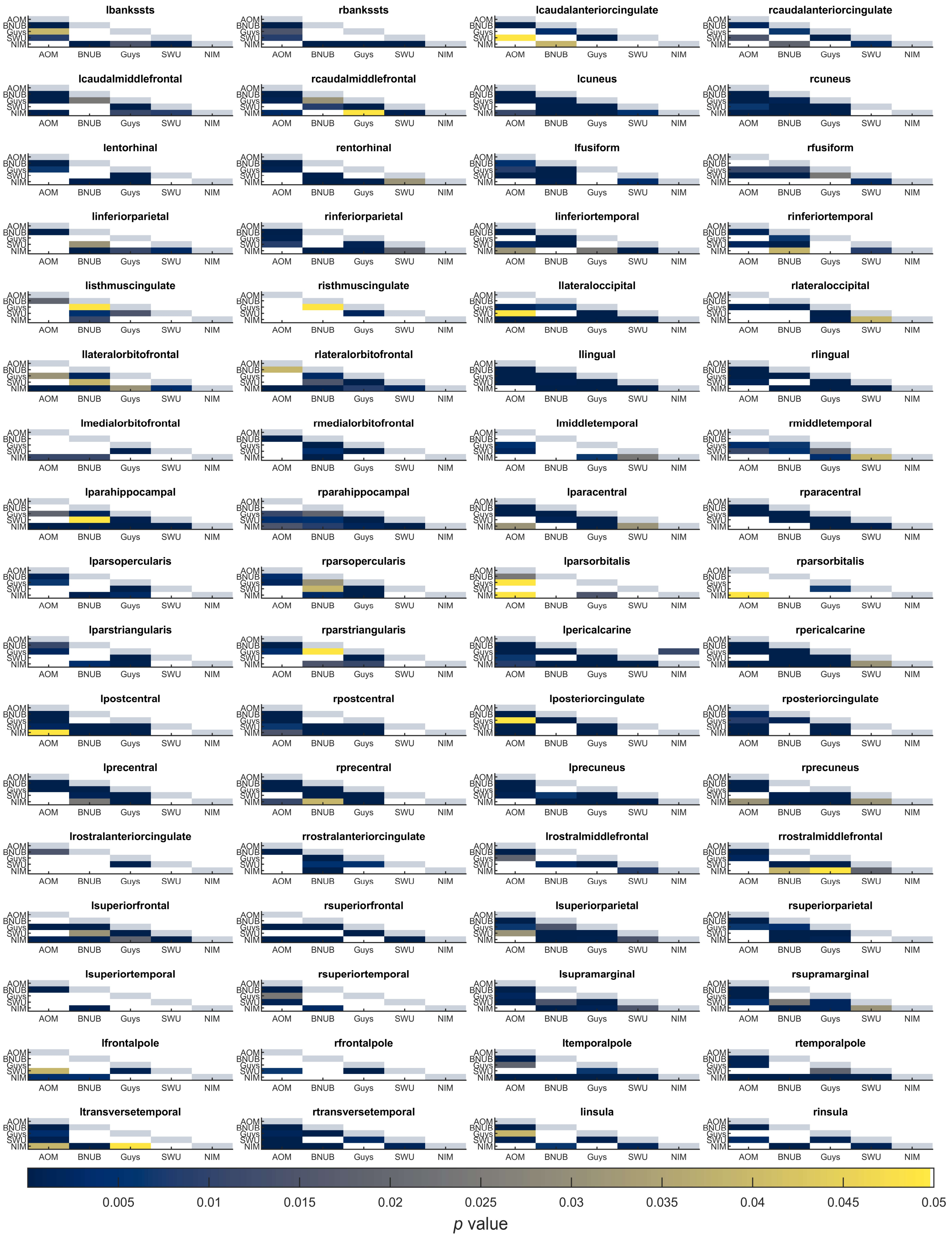


**Figure S1:** Summary of *p*-values from two sample Kolmogorov-Smirnov (KS) test between pairs of scanners for cortical thickness. Each sub-plot indicates the *p*-values before (lower triangle) and after harmonization (upper triangle) between all pairs of scanners; the diagonal elements are shaded in a constant color to help distinguish lower and upper triangles. Each cell is color coded based on their *p*-value and only values smaller than 0.05 are shown. Note that AOMIC dataset has been abbreviated to “AOM”, BNUBeijing dataset has been abbreviated to “BNUB”, and “NIMHANS” dataset has been abbreviated to “NIM”.

### Experiment 1: univariate differences – fractal dimension

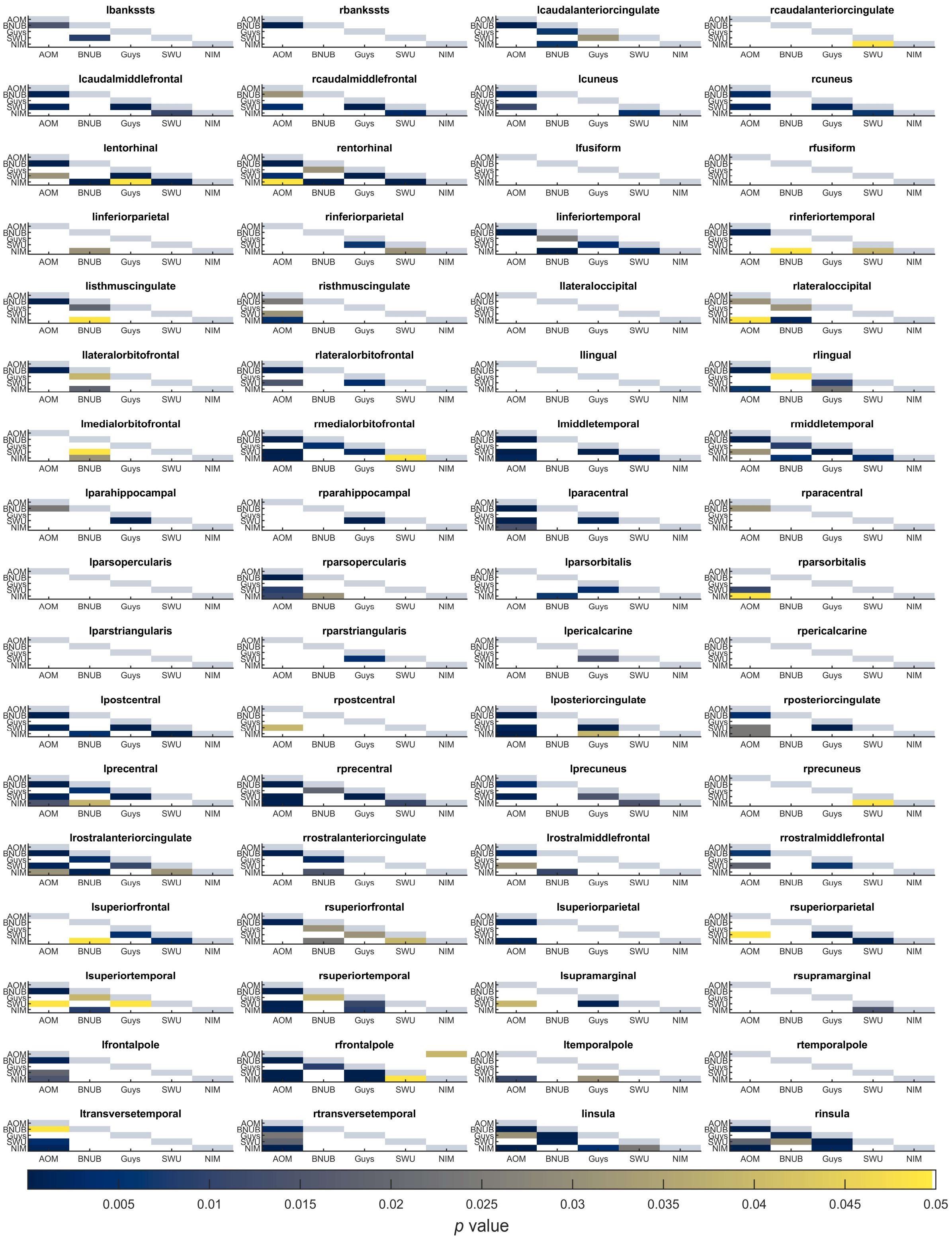

**Figure S2:** Summary of *p*-values from two sample Kolmogorov-Smirnov (KS) test between pairs of scanners for fractal dimension. Each sub-plot indicates the *p*-values before (lower triangle) and after harmonization (upper triangle) between all pairs of scanners; the diagonal elements are shaded in a constant color to help distinguish lower and upper triangles. Each cell is color coded based on their *p*-value and only values smaller than 0.05 are shown. Note that AOMIC dataset has been abbreviated to “AOM”, BNUBeijing dataset has been abbreviated to “BNUB”, and “NIMHANS” dataset has been abbreviated to “NIM”.

### Experiment 1: univariate differences – sulcal depth

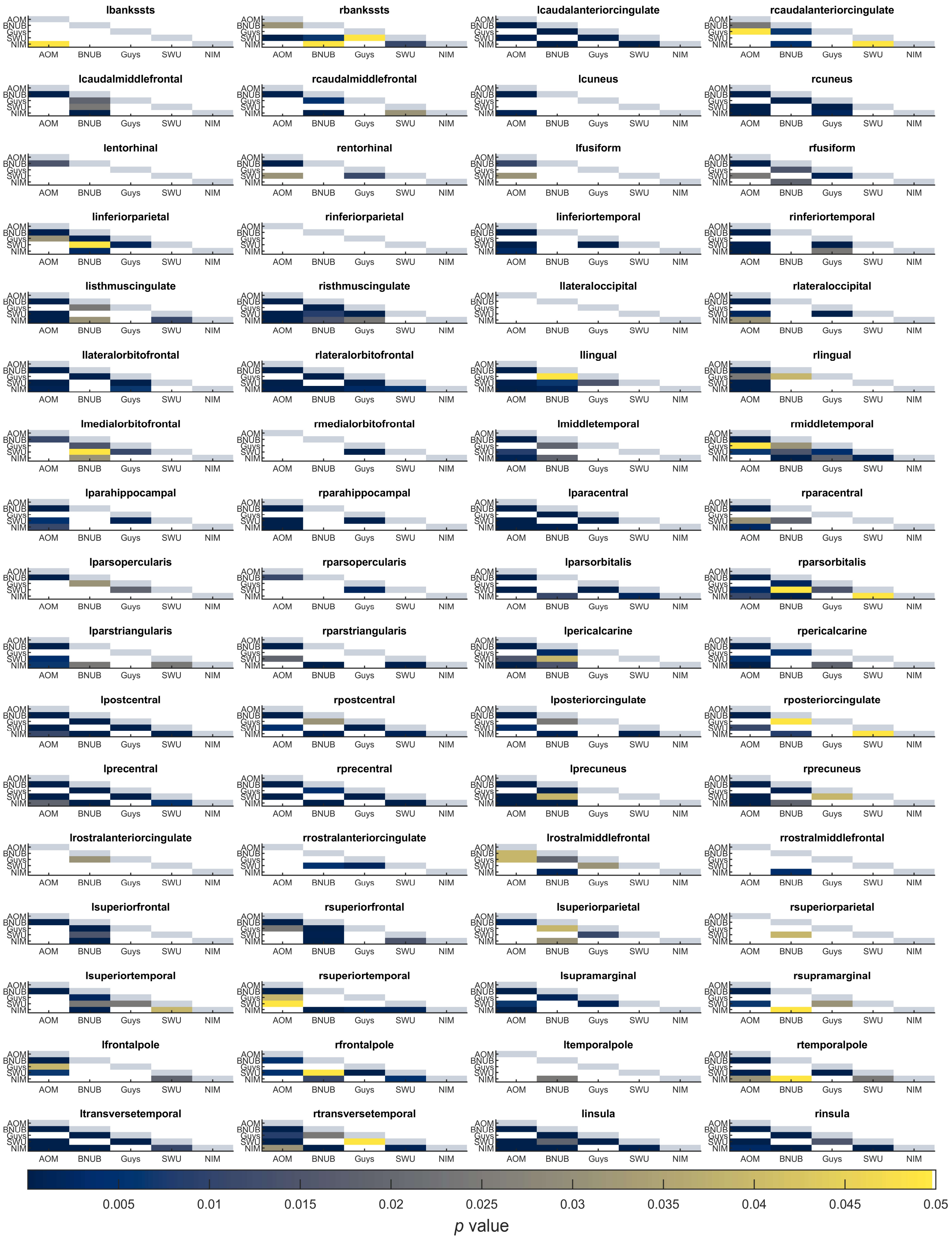


**Figure S3:** Summary of *p*-values from two sample Kolmogorov-Smirnov (KS) test between pairs of scanners for sulcal depth. Each sub-plot indicates the *p*-values before (lower triangle) and after harmonization (upper triangle) between all pairs of scanners; the diagonal elements are shaded in a constant color to help distinguish lower and upper triangles. Each cell is color coded based on their *p*-value and only values smaller than 0.05 are shown. Note that AOMIC dataset has been abbreviated to "AOM", BNUBeijing dataset has been abbreviated to "BNUB", and "NIMHANS" dataset has been abbreviated to "NIM".

### Experiment 1: univariate differences – gyrification index

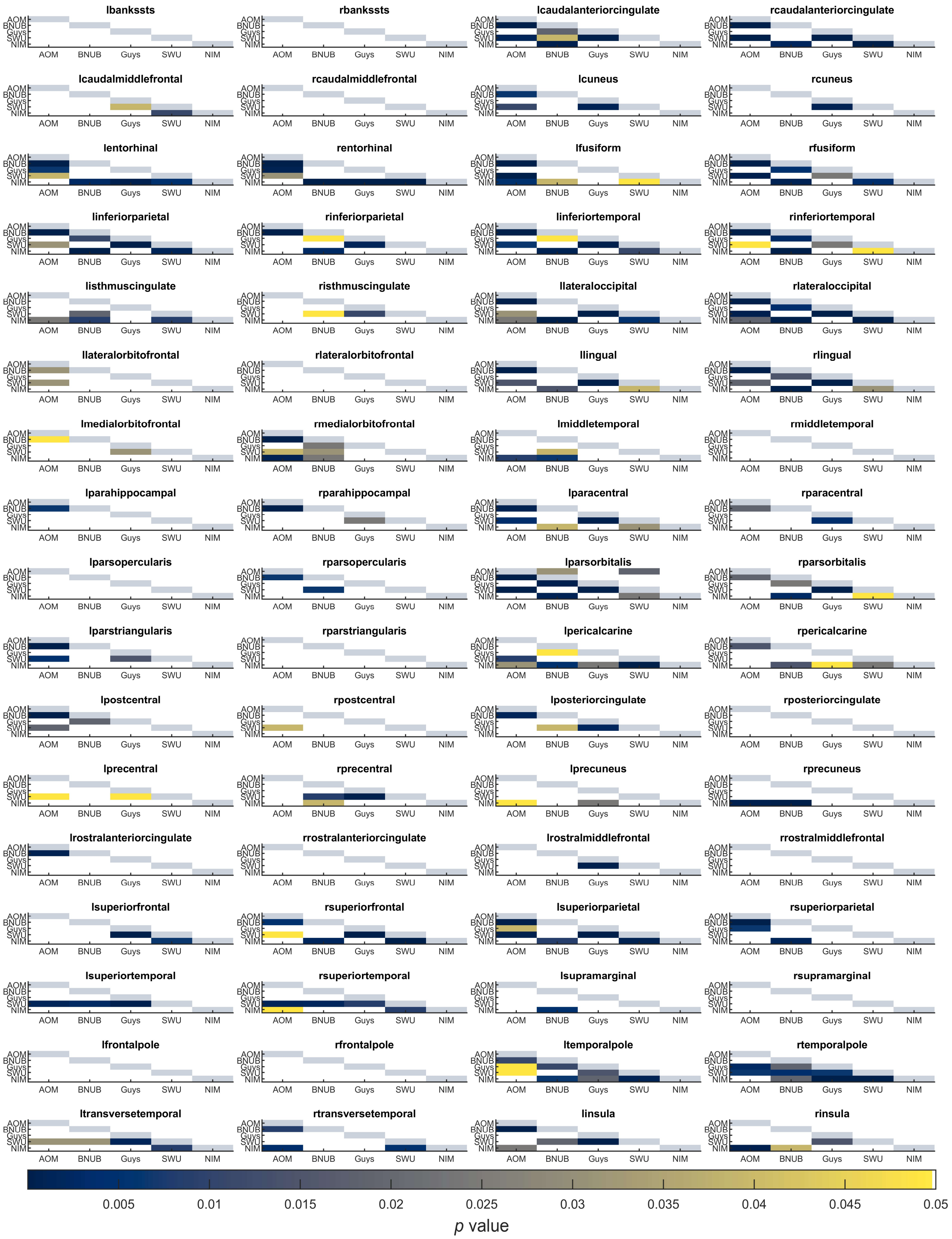


**Figure S4:** Summary of $p$-values from two sample Kolmogorov-Smirnov (KS) test between pairs of scanners for gyrification index. Each sub-plot indicates the $p$-values before (lower triangle) and after harmonization (upper triangle) between all pairs of scanners; the diagonal elements are shaded in a constant color to help distinguish lower and upper triangles. Each cell is color coded based on their $p$-value and only values smaller than 0.05 are shown. Note that AOMIC dataset has been abbreviated to “AOM”, BNUBeijing dataset has been abbreviated to “BNUB”, and “NIMHANS” dataset has been abbreviated to “NIM”.

### Experiment 3: learning curves

#### Volumetric features

##### Three sites

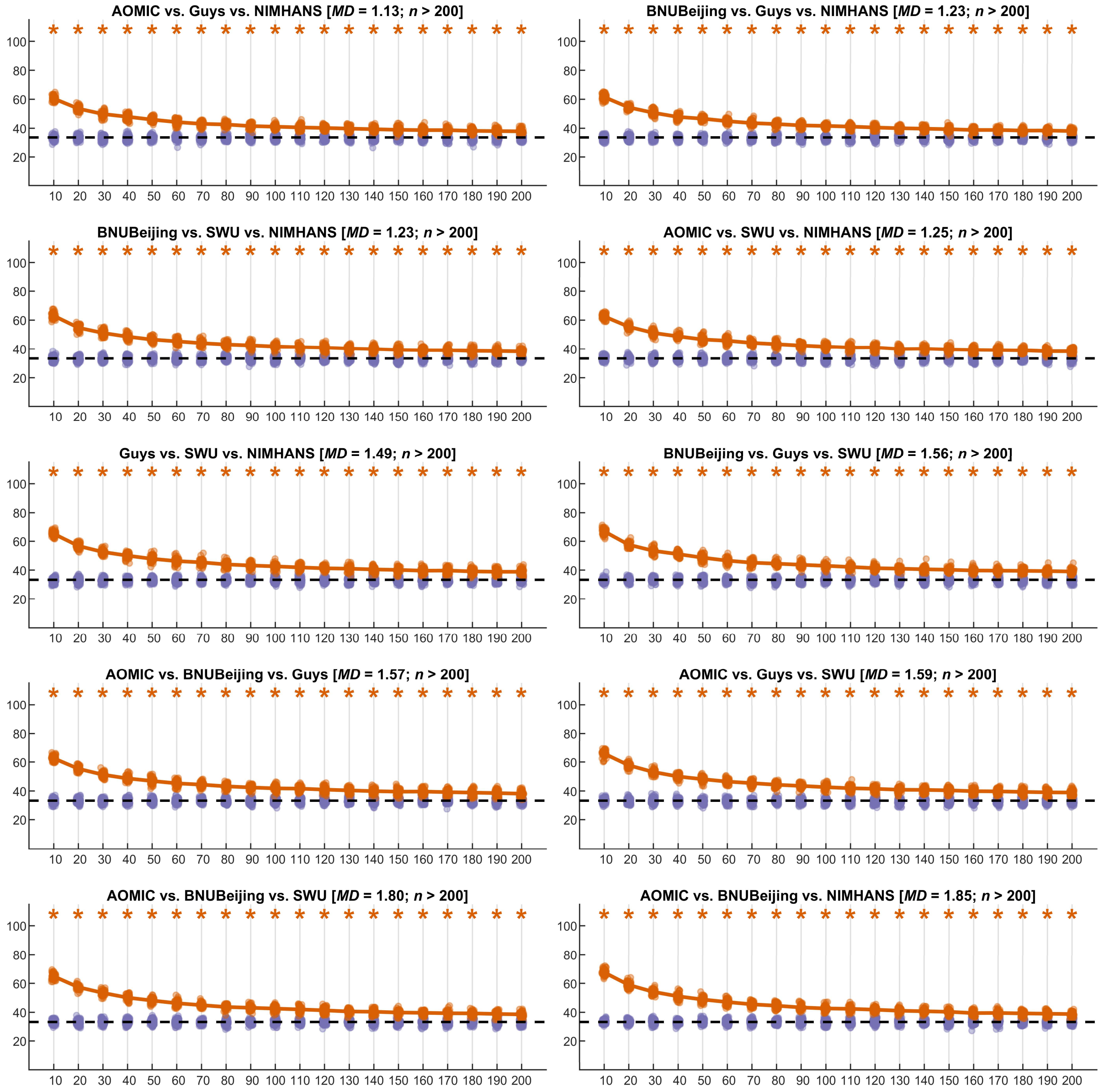


**Figure S5:** Summary of learning curves for volumetric features for three-site combinations; the orange points indicate the test accuracy of the SVM classifier (50 repeats of 10-fold cross-validation), the purple points indicate the permutation test accuracy of the SVM classifier (100 repeats of 10-fold cross-validation), while the dashed black line indicates the theoretical chance accuracy level; the *x*-axis indicates the sample size used for learning harmonization parameters ("*NHLearn*") while the *y*-axis indicates the test accuracy in percentage. The title of each figure indicates the site-combinations, the average Mahalanobis distance (*MD*) of the two sites from the reference, and the sample size required for learning harmonization parameter (*n*) such that the SVM classifier performance was no different than chance level; the accuracies that were above chance are marked with an orange asterisk mark

#### Four and five sites

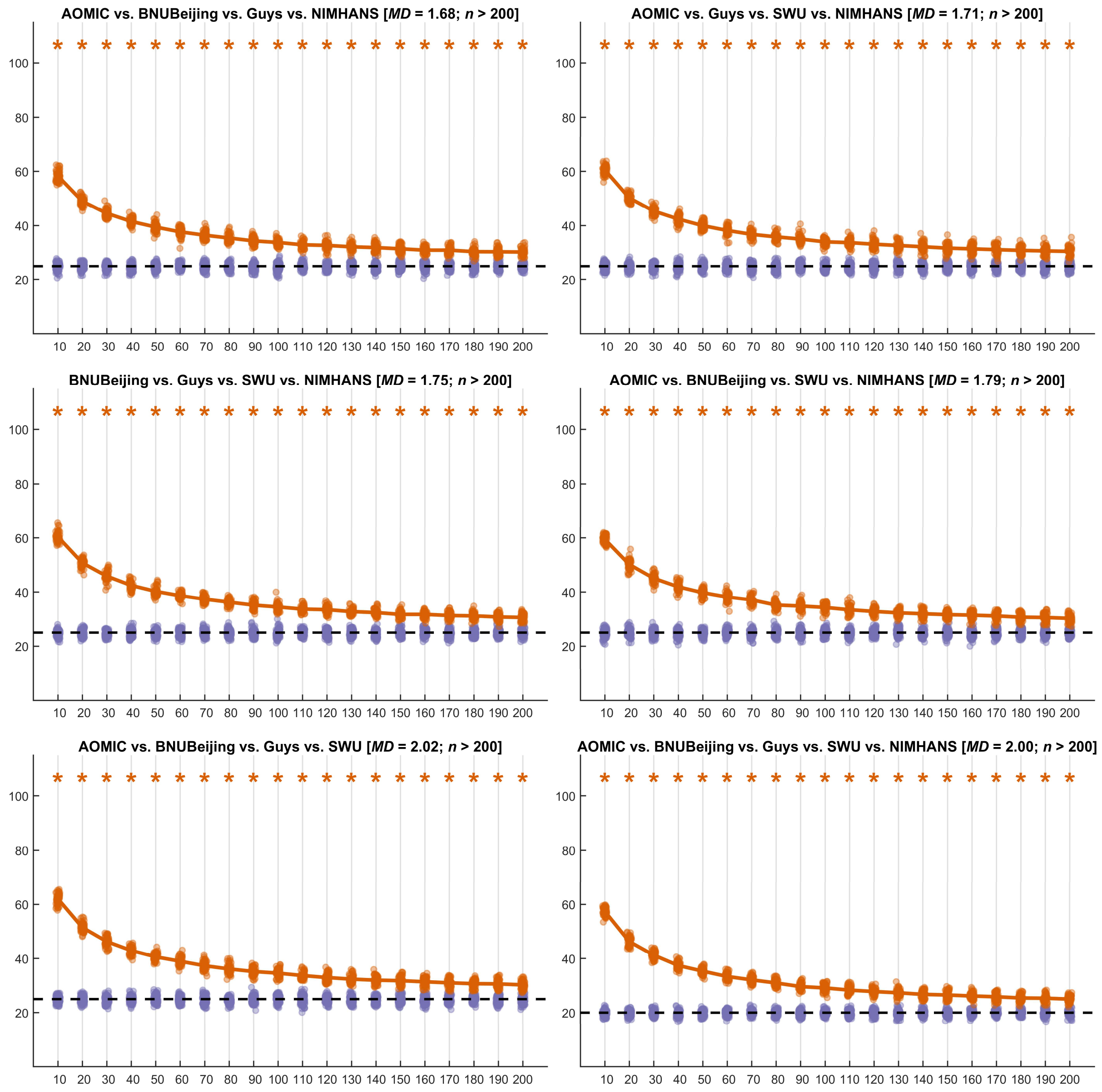


**Figure S6:** Summary of learning curves for volumetric features for four-site and five-site combinations; the orange points indicate the test accuracy of the SVM classifier (50 repeats of 10-fold cross-validation), the purple points indicate the permutation test accuracy of the SVM classifier (100 repeats of 10-fold cross-validation), while the dashed black line indicates the theoretical chance accuracy level; the *x*-axis indicates the sample size used for learning harmonization parameters ("*NHLearn*") while the *y*-axis indicates the test accuracy in percentage. The title of each figure indicates the site-combinations, the average Mahalanobis distance (*MD*) of the two sites from the reference, and the sample size required for learning harmonization parameter (*n*) such that the SVM classifier performance was no different than chance level; the accuracies that were above chance are marked with an orange asterisk mark

#### Cortical thickness

##### Two sites

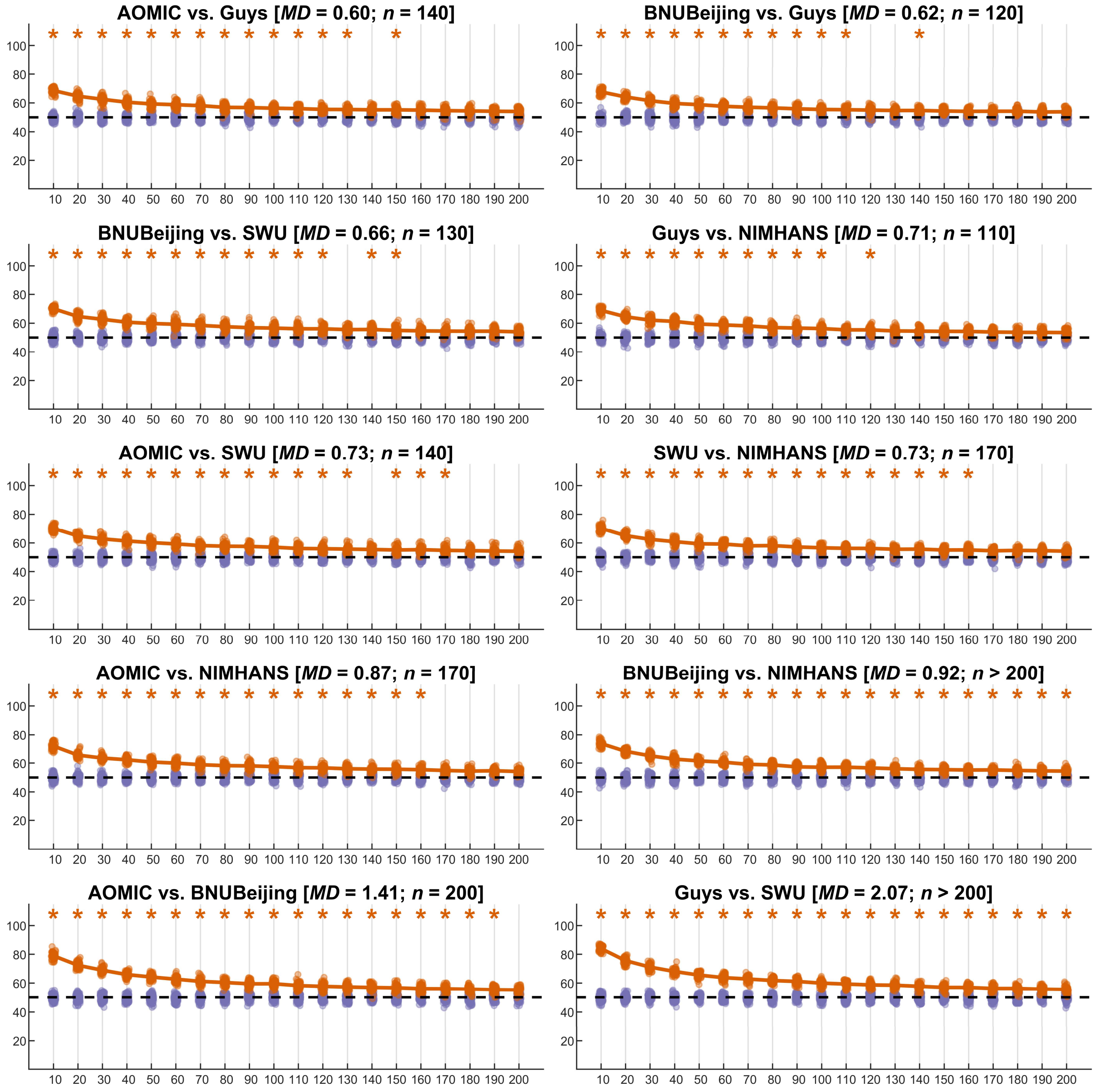


**Figure S7:** Summary of learning curves for cortical thickness features for two-site combinations; the orange points indicate the test accuracy of the SVM classifier (50 repeats of 10-fold cross-validation), the purple points indicate the permutation test accuracy of the SVM classifier (100 repeats of 10-fold cross-validation), while the dashed black line indicates the theoretical chance accuracy level; the *x*-axis indicates the sample size used for learning harmonization parameters ("*NHLearn*") while the *y*-axis indicates the test accuracy in percentage. The title of each figure indicates the site-combinations, the average Mahalanobis distance (*MD*) of the two sites from the reference, and the sample size required for learning harmonization parameter (*n*) such that the SVM classifier performance was no different than chance level; the accuracies that were above chance are marked with an orange asterisk mark

#### Three sites

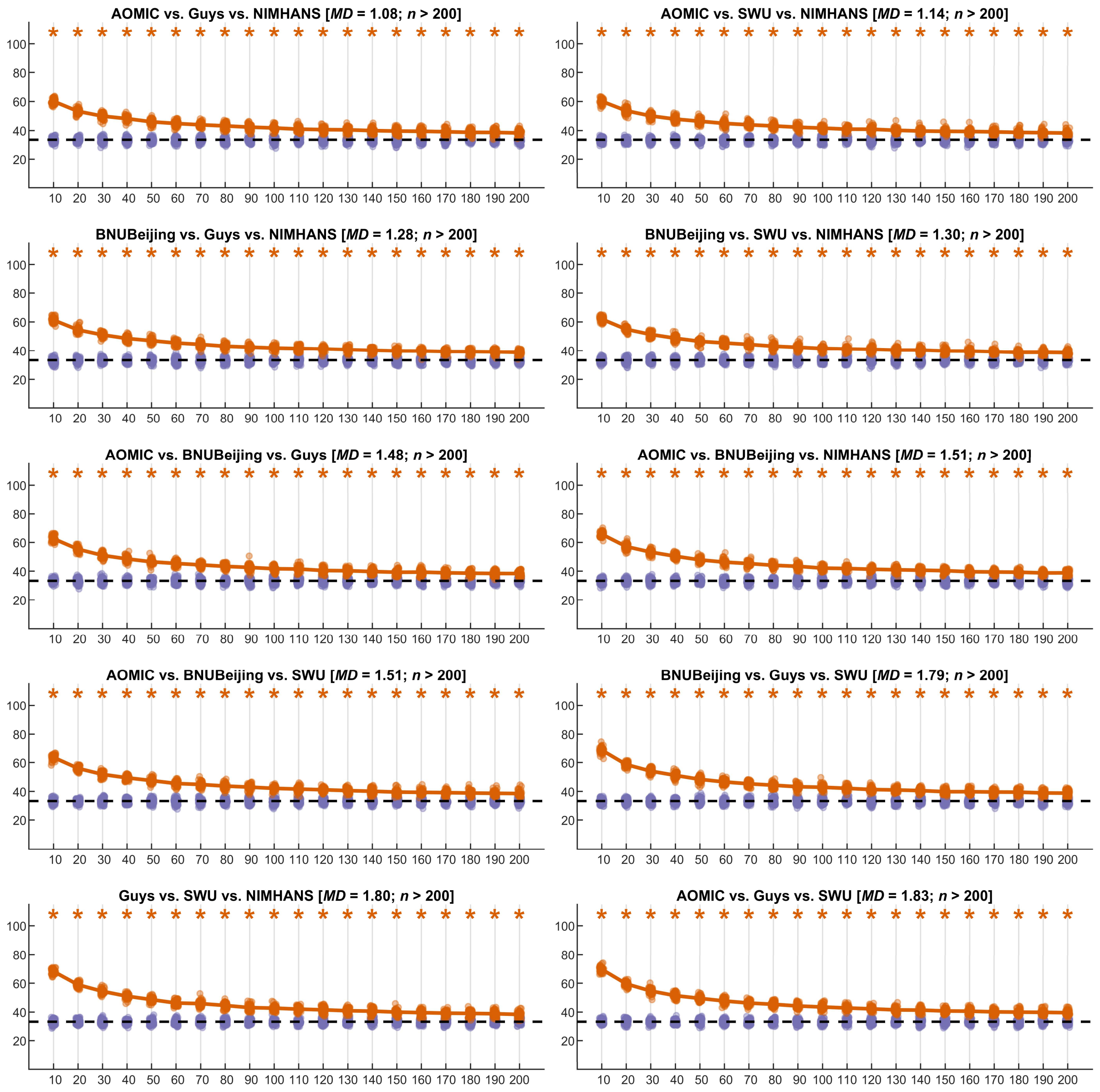


**Figure S8:** Summary of learning curves for cortical thickness features for three-site combinations; the orange points indicate the test accuracy of the SVM classifier (50 repeats of 10-fold cross-validation), the purple points indicate the permutation test accuracy of the SVM classifier (100 repeats of 10-fold cross-validation), while the dashed black line indicates the theoretical chance accuracy level; the *x*-axis indicates the sample size used for learning harmonization parameters ("*NHLearn*") while the *y*-axis indicates the test accuracy in percentage. The title of each figure indicates the site-combinations, the average Mahalanobis distance (*MD*) of the two sites from the reference, and the sample size required for learning harmonization parameter (*n*) such that the SVM classifier performance was no different than chance level; the accuracies that were above chance are marked with an orange asterisk mark

#### Four and five sites

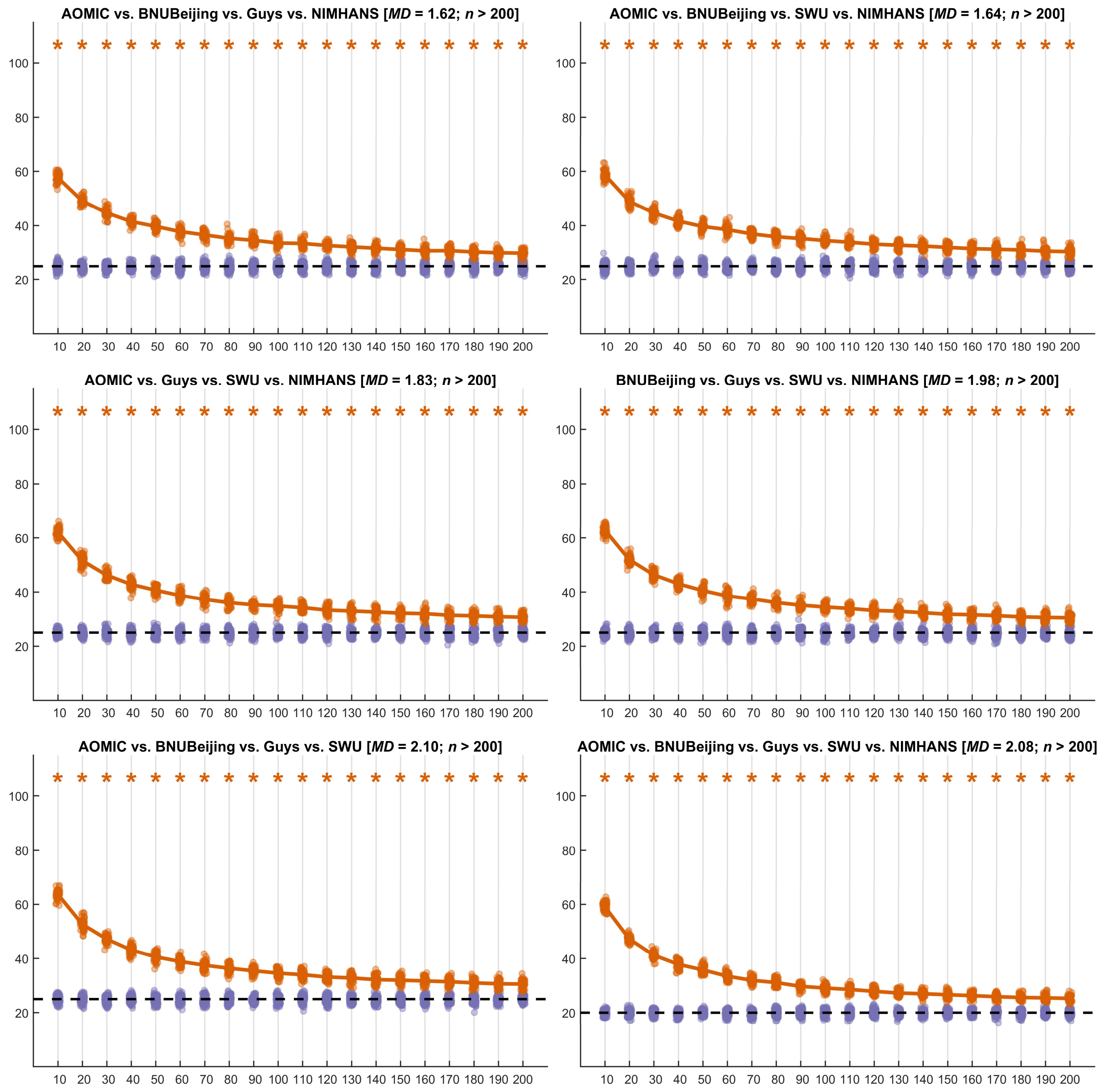


**Figure S9:** Summary of learning curves for cortical thickness features for four-site and five-site combinations; the orange points indicate the test accuracy of the SVM classifier (50 repeats of 10-fold cross-validation), the purple points indicate the permutation test accuracy of the SVM classifier (100 repeats of 10-fold cross-validation), while the dashed black line indicates the theoretical chance accuracy level; the *x*-axis indicates the sample size used for learning harmonization parameters ("*NHLearn*") while the *y*-axis indicates the test accuracy in percentage. The title of each figure indicates the site-combinations, the average Mahalanobis distance (*MD*) of the two sites from the reference, and the sample size required for learning harmonization parameter (*n*) such that the SVM classifier performance was no different than chance level; the accuracies that were above chance are marked with an orange asterisk mark

#### Fractal dimension

##### Two sites

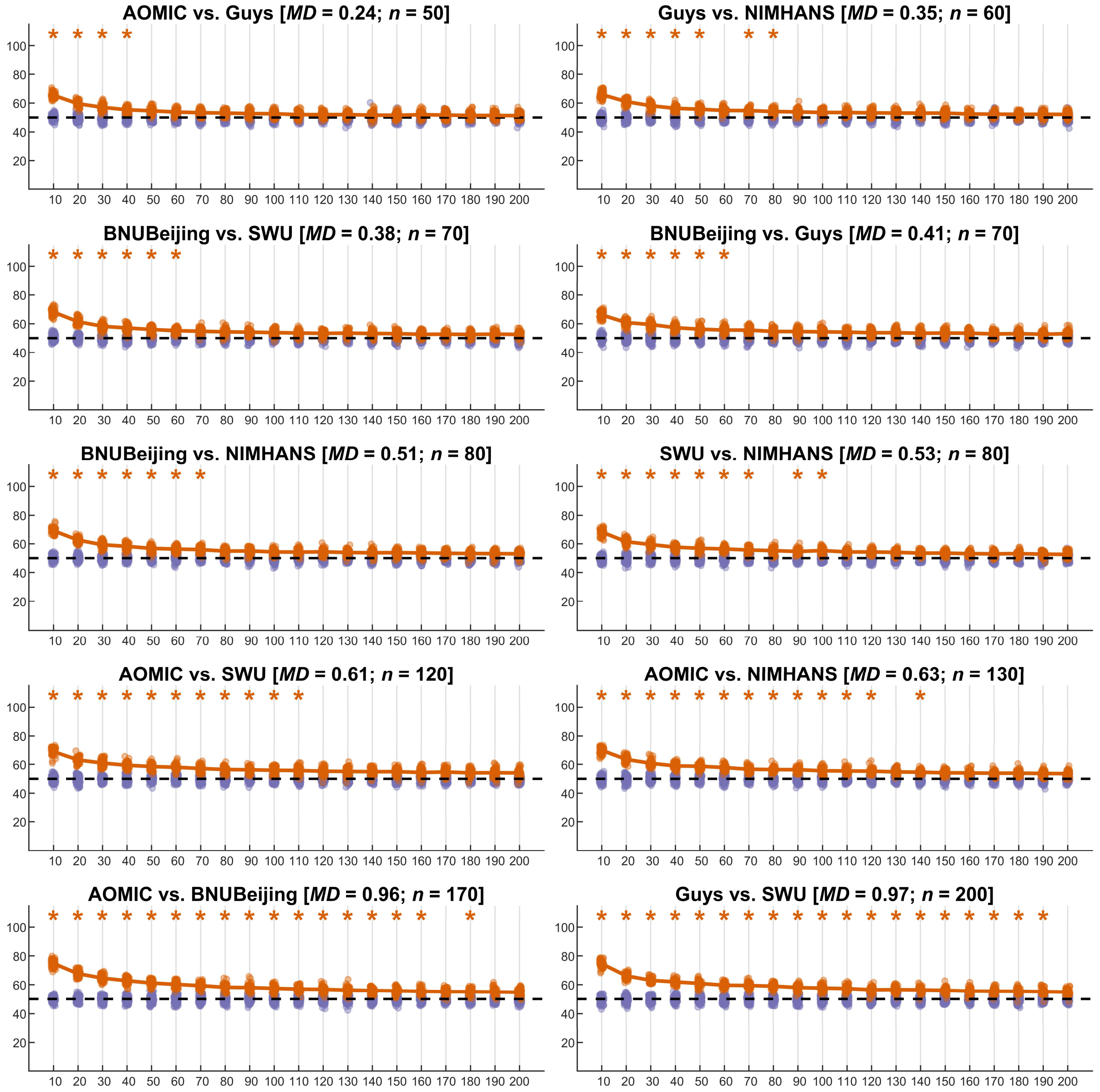


**Figure S10:** Summary of learning curves for fractal dimension features for two-site combinations; the orange points indicate the test accuracy of the SVM classifier (50 repeats of 10-fold cross-validation), the purple points indicate the permutation test accuracy of the SVM classifier (100 repeats of 10-fold cross-validation), while the dashed black line indicates the theoretical chance accuracy level; the *x*-axis indicates the sample size used for learning harmonization parameters ("*NHLearn*") while the *y*-axis indicates the test accuracy in percentage. The title of each figure indicates the site-combinations, the average Mahalanobis distance (*MD*) of the two sites from the reference, and the sample size required for learning harmonization parameter (*n*) such that the SVM classifier performance was no different than chance level; the accuracies that were above chance are marked with an orange asterisk mark

#### Three sites

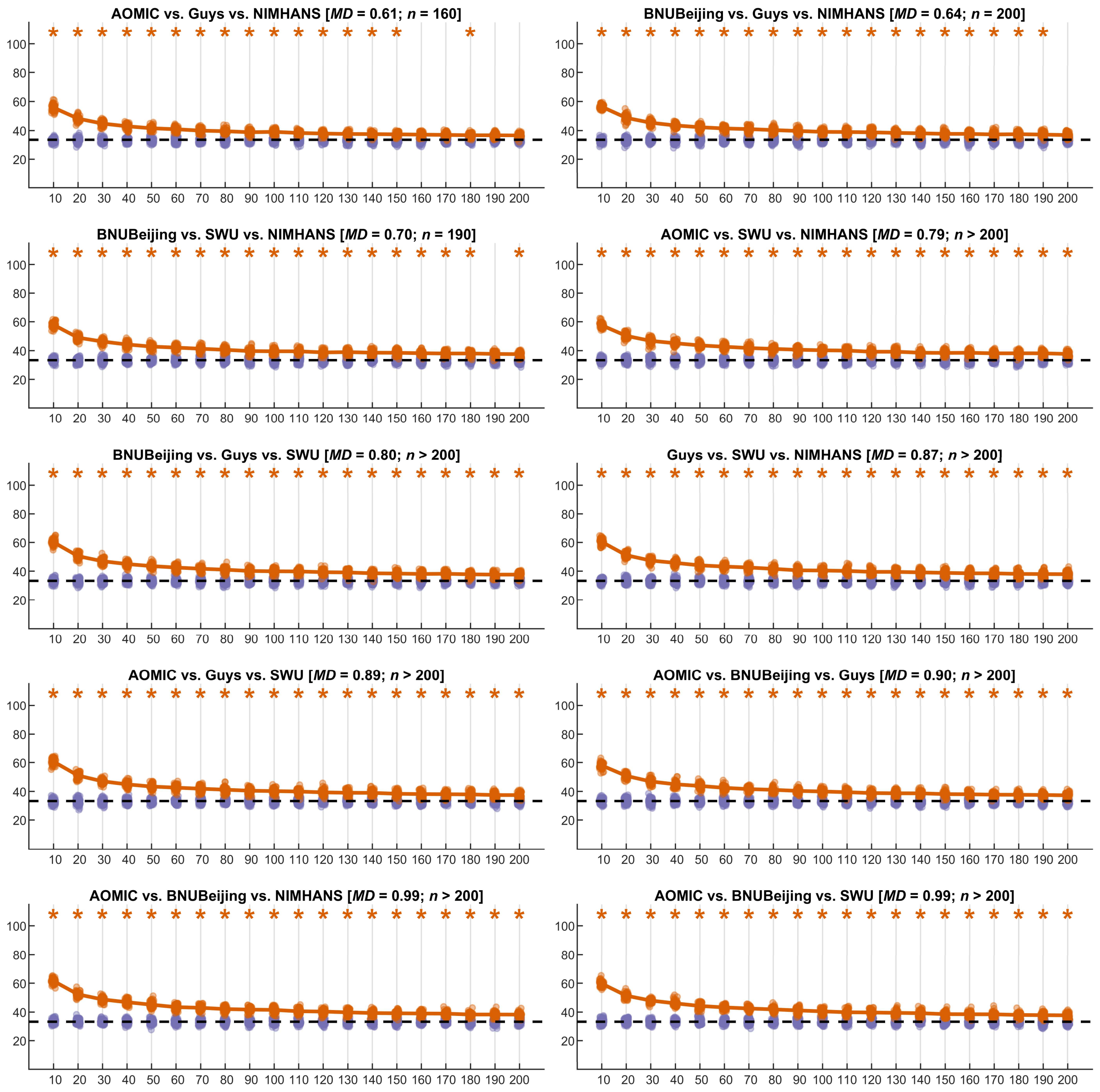


**Figure S11:** Summary of learning curves for fractal dimension features for three-site combinations; the orange points indicate the test accuracy of the SVM classifier (50 repeats of 10-fold cross-validation), the purple points indicate the permutation test accuracy of the SVM classifier (100 repeats of 10-fold cross-validation), while the dashed black line indicates the theoretical chance accuracy level; the *x*-axis indicates the sample size used for learning harmonization parameters ("*NHLearn*") while the *y*-axis indicates the test accuracy in percentage. The title of each figure indicates the site-combinations, the average Mahalanobis distance (*MD*) of the two sites from the reference, and the sample size required for learning harmonization parameter (*n*) such that the SVM classifier performance was no different than chance level; the accuracies that were above chance are marked with an orange asterisk mark

#### Four and five sites

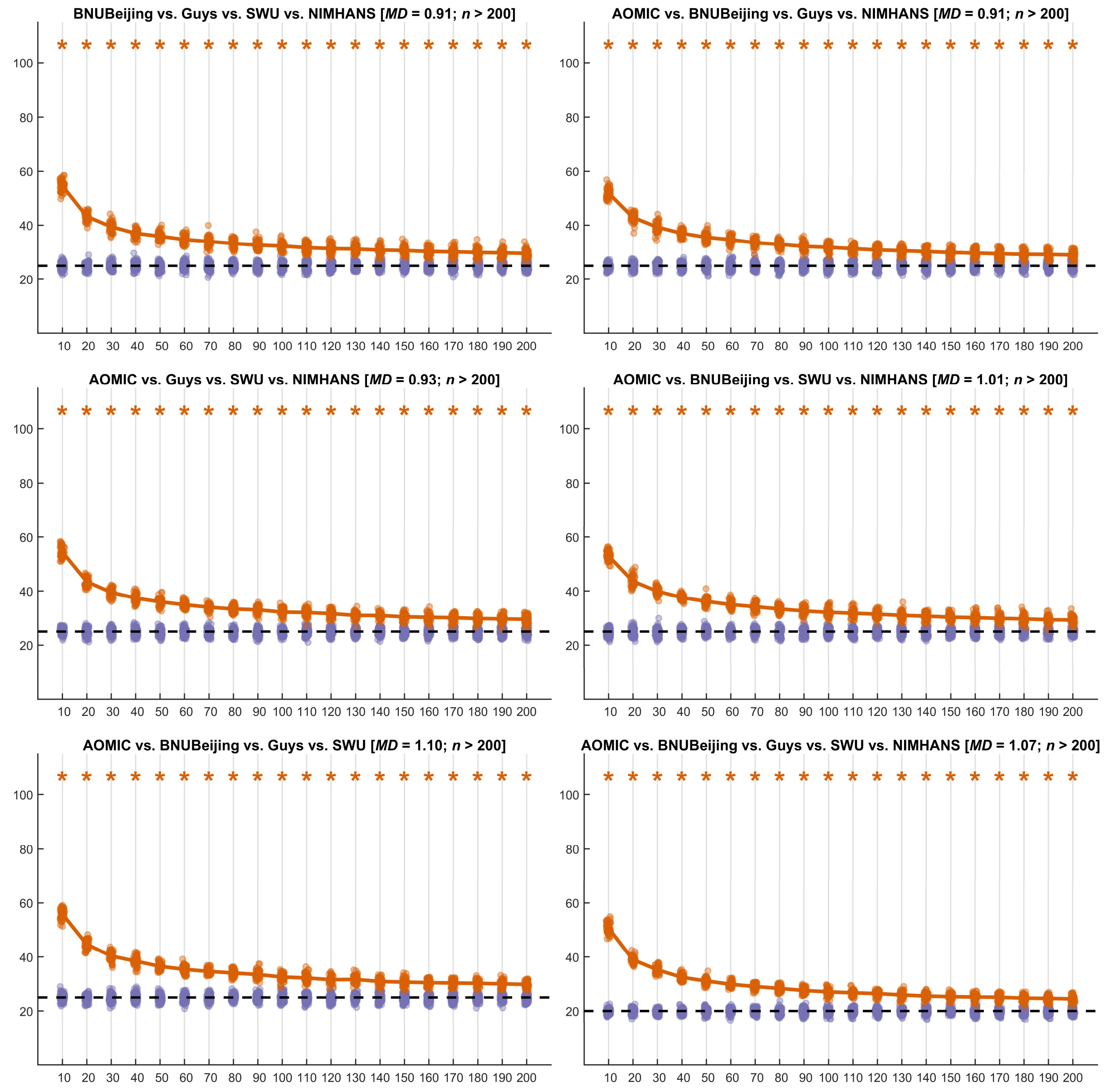


**Figure S12:** Summary of learning curves for fractal dimension features for four-site and five-site combinations; the orange points indicate the test accuracy of the SVM classifier (50 repeats of 10-fold cross-validation), the purple points indicate the permutation test accuracy of the SVM classifier (100 repeats of 10-fold cross-validation), while the dashed black line indicates the theoretical chance accuracy level; the *x*-axis indicates the sample size used for learning harmonization parameters ("*NHLearn*") while the *y*-axis indicates the test accuracy in percentage. The title of each figure indicates the site-combinations, the average Mahalanobis distance (*MD*) of the two sites from the reference, and the sample size required for learning harmonization parameter (*n*) such that the SVM classifier performance was no different than chance level; the accuracies that were above chance are marked with an orange asterisk mark

Sulcal depth

#### Two sites

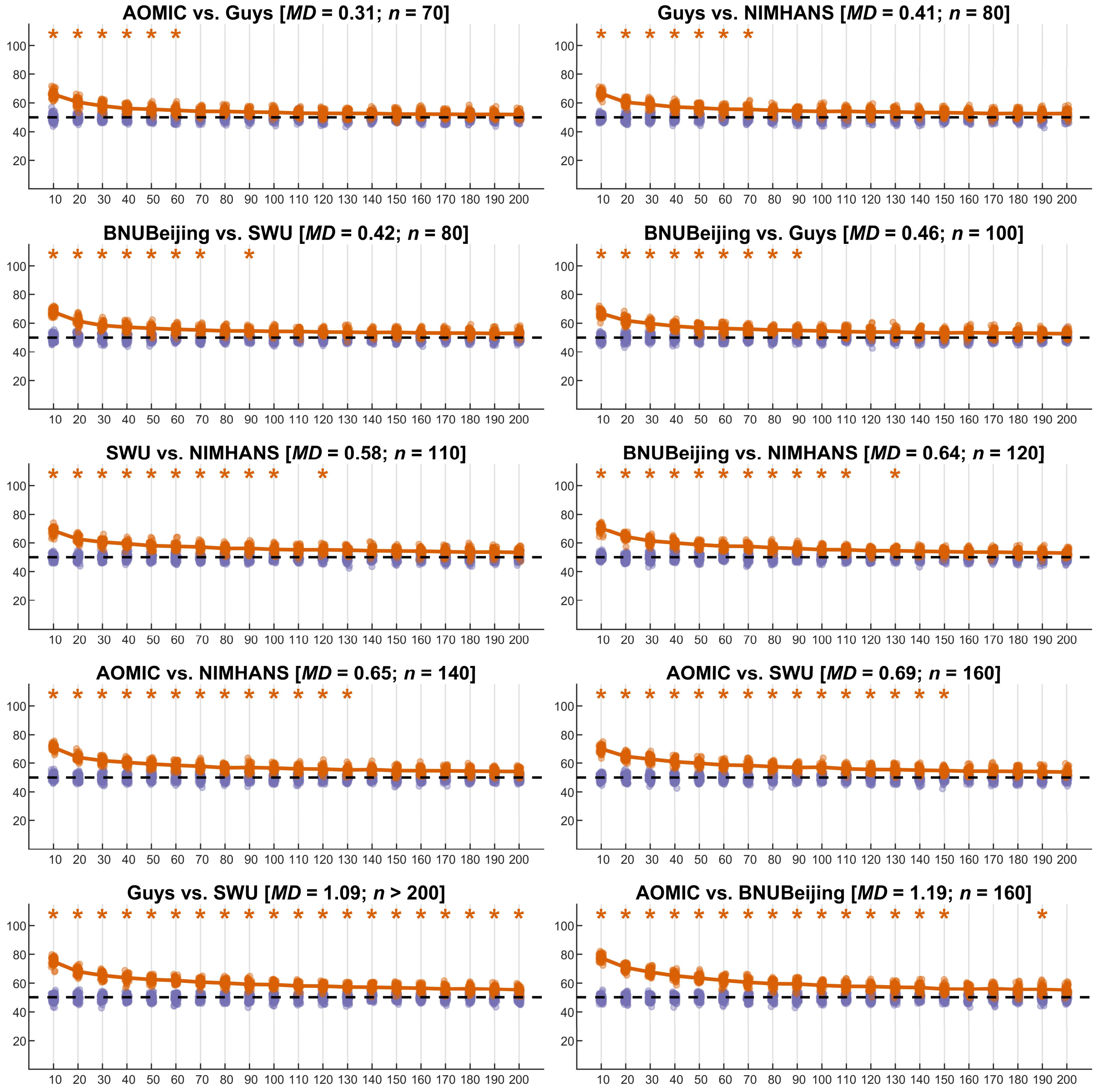


**Figure S13:** Summary of learning curves for sulcal depth features for two-site combinations; the orange points indicate the test accuracy of the SVM classifier (50 repeats of 10-fold cross-validation), the purple points indicate the permutation test accuracy of the SVM classifier (100 repeats of 10-fold cross-validation), while the dashed black line indicates the theoretical chance accuracy level; the *x*-axis indicates the sample size used for learning harmonization parameters ("*NHLearn*") while the *y*-axis indicates the test accuracy in percentage. The title of each figure indicates the site-combinations, the average Mahalanobis distance (*MD*) of the two sites from the reference, and the sample size required for learning harmonization parameter (*n*) such that the SVM classifier performance was no different than chance level; the accuracies that were above chance are marked with an orange asterisk mark

#### Three sites

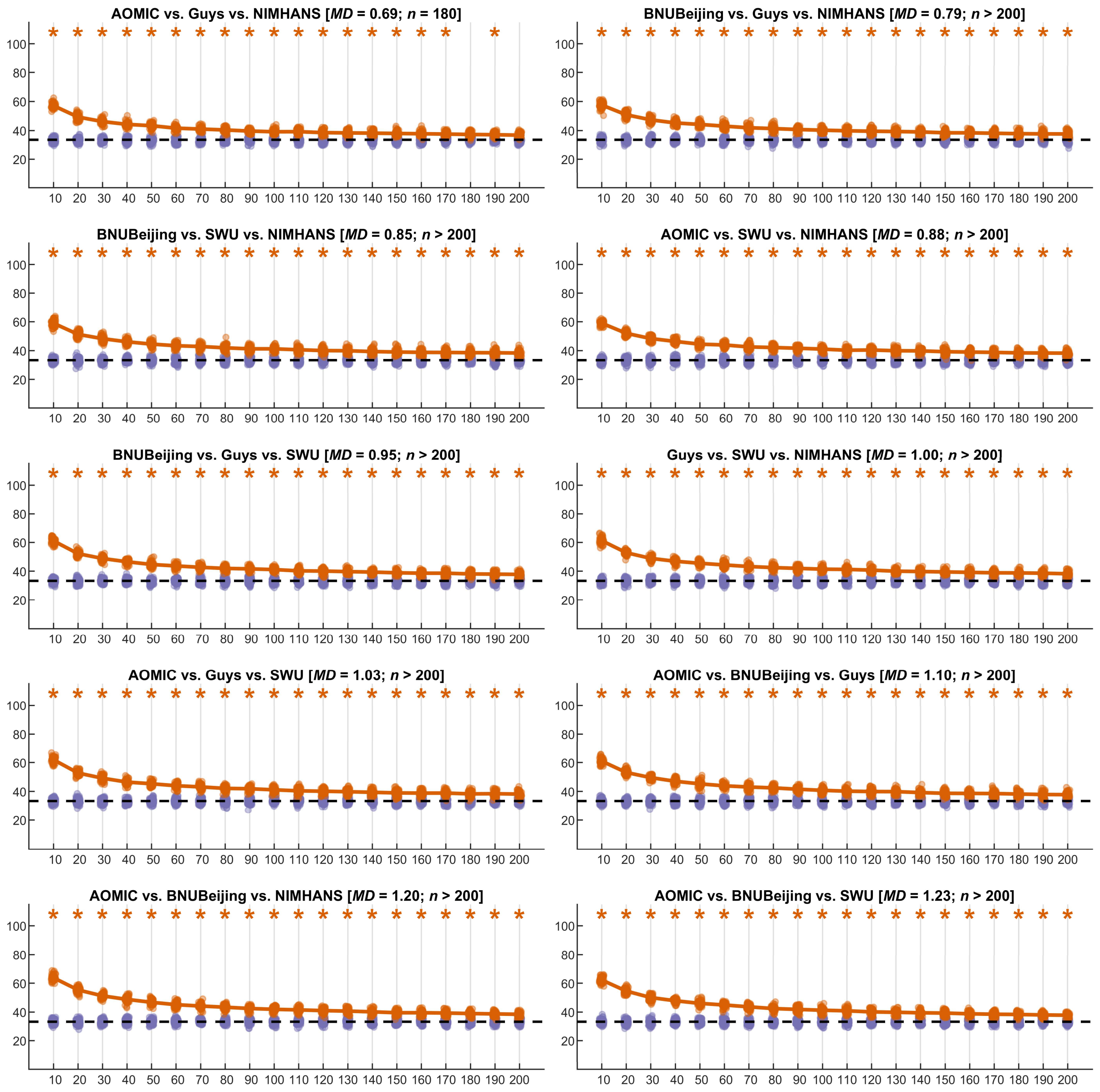


**Figure S14:** Summary of learning curves for sulcal depth features for three-site combinations; the orange points indicate the test accuracy of the SVM classifier (50 repeats of 10-fold cross-validation), the purple points indicate the permutation test accuracy of the SVM classifier (100 repeats of 10-fold cross-validation), while the dashed black line indicates the theoretical chance accuracy level; the $x$-axis indicates the sample size used for learning harmonization parameters ("*NHLearn*") while the $y$-axis indicates the test accuracy in percentage. The title of each figure indicates the site-combinations, the average Mahalanobis distance (*MD*) of the two sites from the reference, and the sample size required for learning harmonization parameter ($n$) such that the SVM classifier performance was no different than chance level; the accuracies that were above chance are marked with an orange asterisk mark

#### Four and five sites

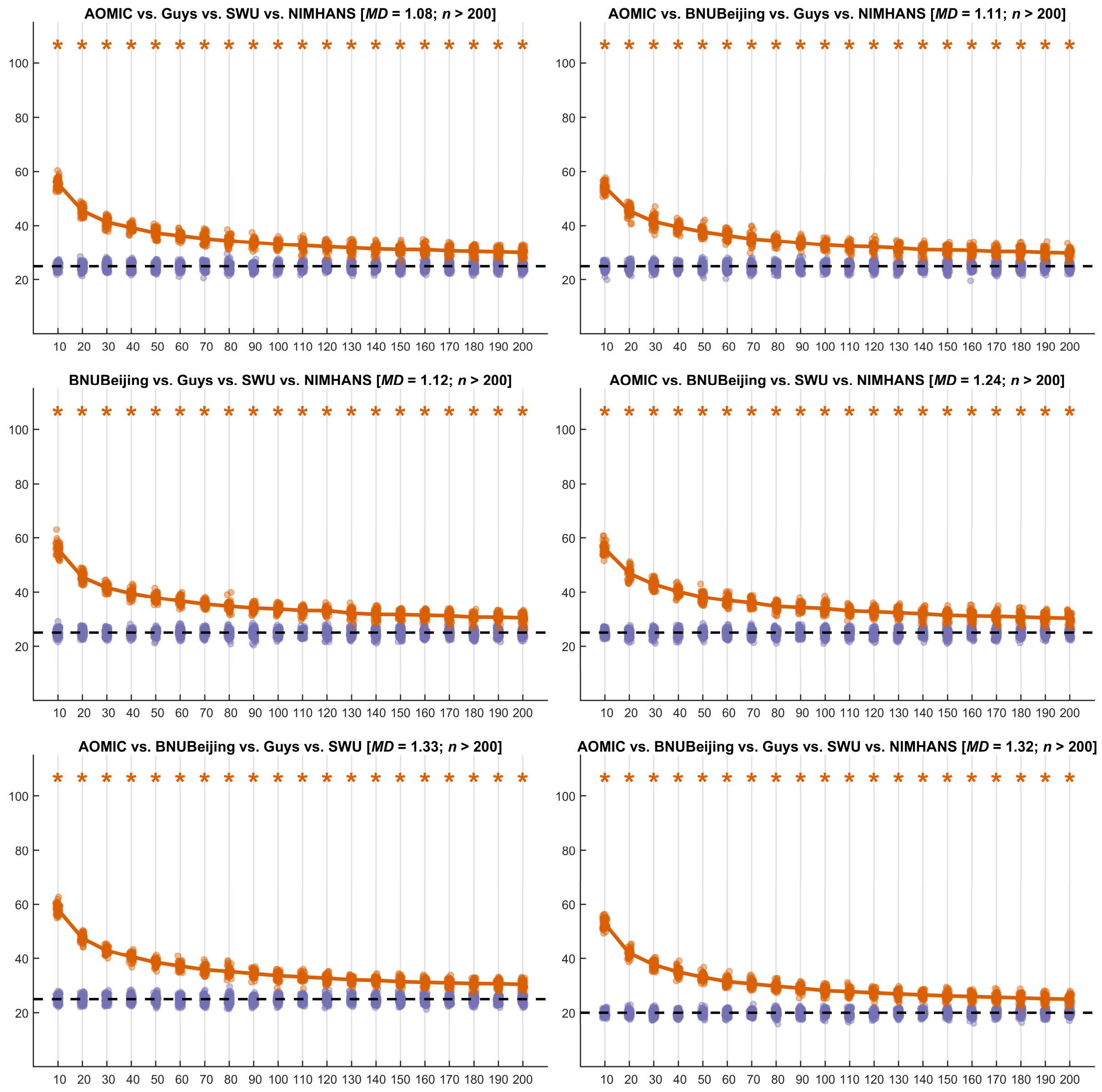


**Figure S15:** Summary of learning curves for sulcal depth features for four-site and five-site combinations; the orange points indicate the test accuracy of the SVM classifier (50 repeats of 10-fold cross-validation), the purple points indicate the permutation test accuracy of the SVM classifier (100 repeats of 10-fold cross-validation), while the dashed black line indicates the theoretical chance accuracy level; the $x$-axis indicates the sample size used for learning harmonization parameters ("*NHLearn*") while the $y$-axis indicates the test accuracy in percentage. The title of each figure indicates the site-combinations, the average Mahalanobis distance (*MD*) of the two sites from the reference, and the sample size required for learning harmonization parameter ($n$) such that the SVM classifier performance was no different than chance level; the accuracies that were above chance are marked with an orange asterisk mark

### Gyrification index

#### Two sites

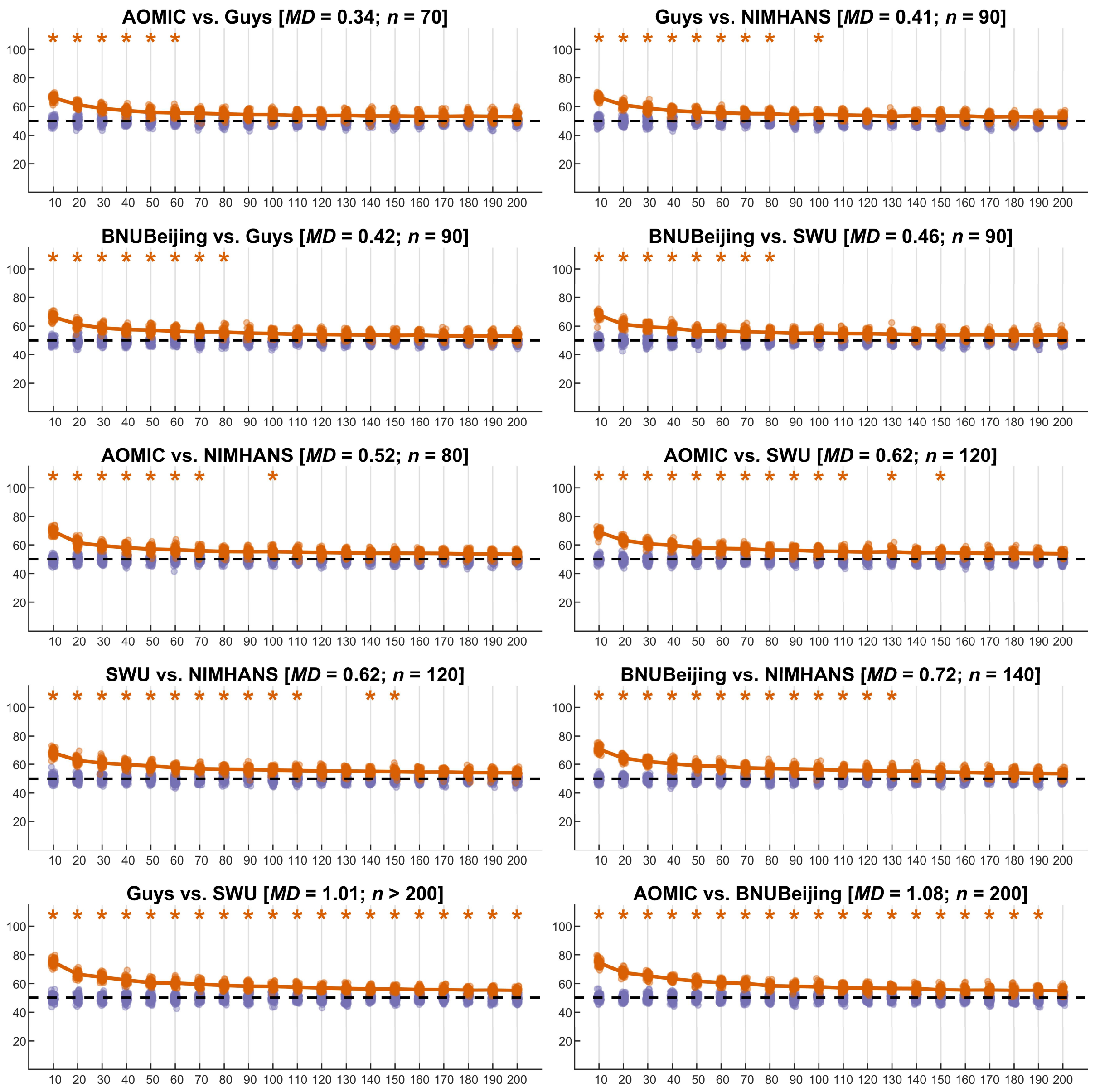


**Figure S16:** Summary of learning curves for gyrification index features for two-site combinations; the orange points indicate the test accuracy of the SVM classifier (50 repeats of 10-fold cross-validation), the purple points indicate the permutation test accuracy of the SVM classifier (100 repeats of 10-fold cross-validation), while the dashed black line indicates the theoretical chance accuracy level; the $x$-axis indicates the sample size used for learning harmonization parameters ("*NHLearn*") while the $y$-axis indicates the test accuracy in percentage. The title of each figure indicates the site-combinations, the average Mahalanobis distance (*MD*) of the two sites from the reference, and the sample size required for learning harmonization parameter ($n$) such that the SVM classifier performance was no different than chance level; the accuracies that were above chance are marked with an orange asterisk mark

#### Three sites

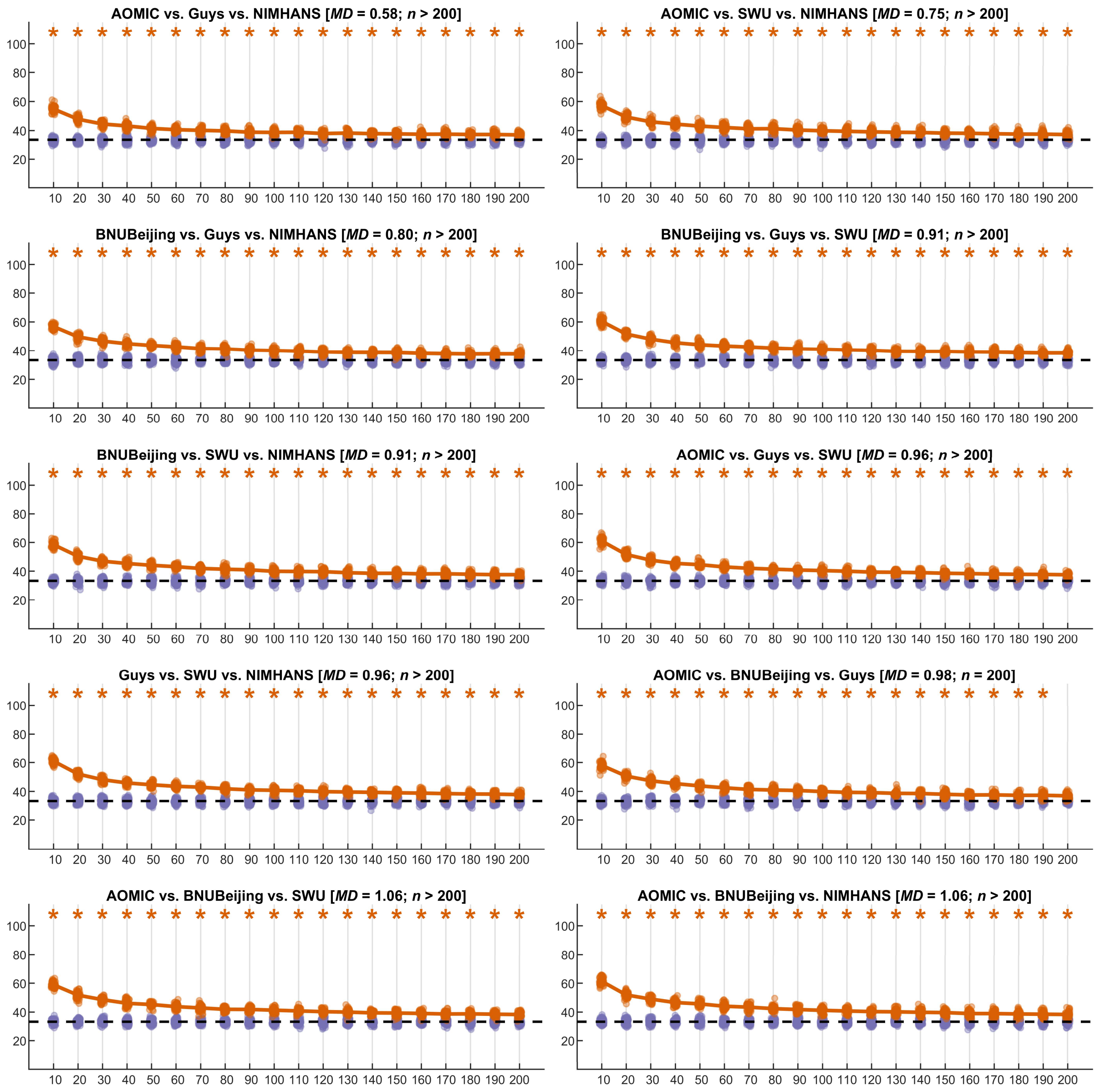


**Figure S17:** Summary of learning curves for gyrification index features for three-site combinations; the orange points indicate the test accuracy of the SVM classifier (50 repeats of 10-fold cross-validation), the purple points indicate the permutation test accuracy of the SVM classifier (100 repeats of 10-fold cross-validation), while the dashed black line indicates the theoretical chance accuracy level; the *x*-axis indicates the sample size used for learning harmonization parameters ("*NHLearn*") while the *y*-axis indicates the test accuracy in percentage. The title of each figure indicates the site-combinations, the average Mahalanobis distance (*MD*) of the two sites from the reference, and the sample size required for learning harmonization parameter (*n*) such that the SVM classifier performance was no different than chance level; the accuracies that were above chance are marked with an orange asterisk mark

#### Four and five sites

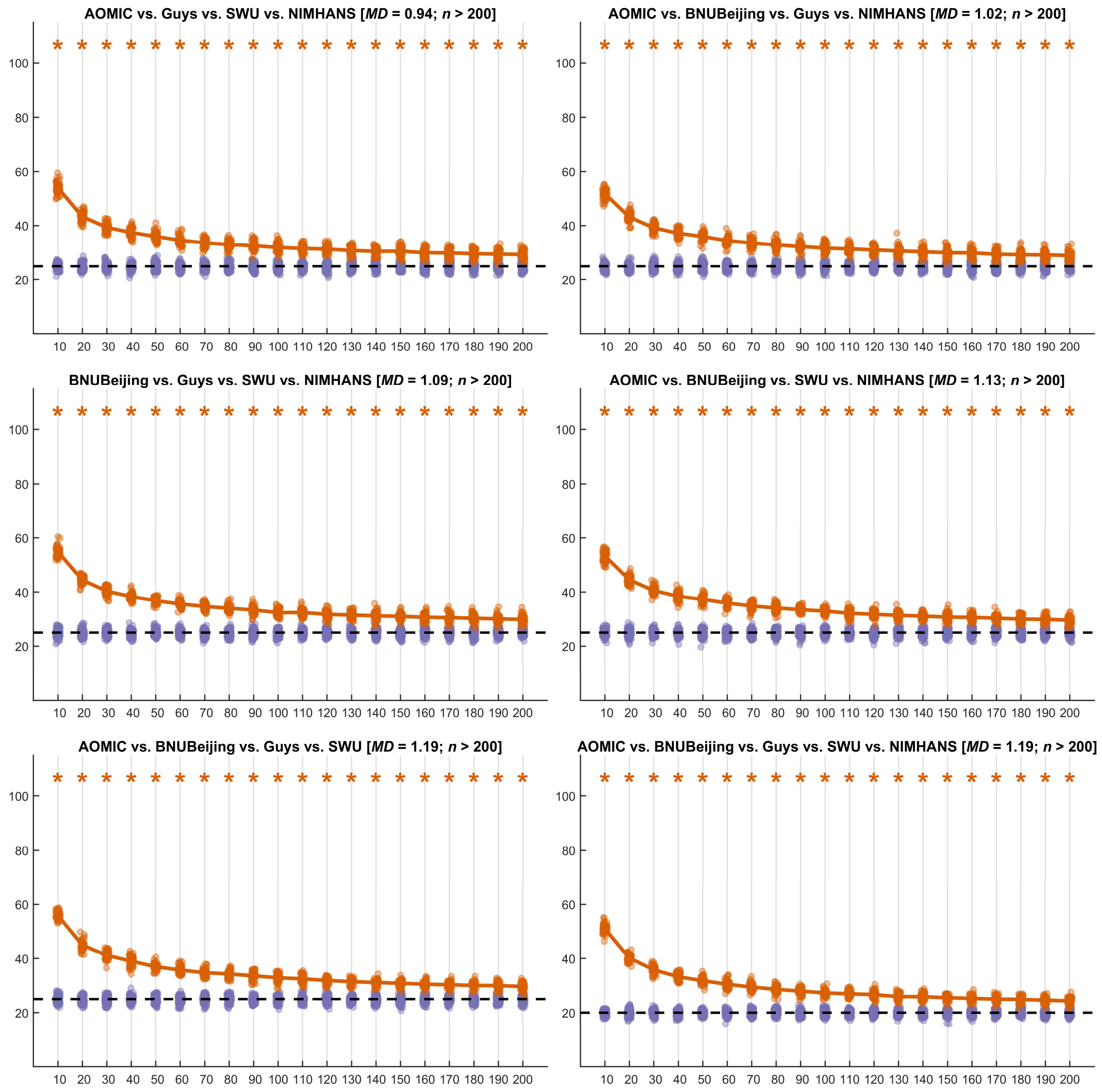


**Figure S18:** Summary of learning curves for gyrification index features for four-site and five-site combinations; the orange points indicate the test accuracy of the SVM classifier (50 repeats of 10-fold cross-validation), the purple points indicate the permutation test accuracy of the SVM classifier (100 repeats of 10-fold cross-validation), while the dashed black line indicates the theoretical chance accuracy level; the *x*-axis indicates the sample size used for learning harmonization parameters ("*NHLearn*") while the *y*-axis indicates the test accuracy in percentage. The title of each figure indicates the site-combinations, the average Mahalanobis distance (*MD*) of the two sites from the reference, and the sample size required for learning harmonization parameter (*n*) such that the SVM classifier performance was no different than chance level; the accuracies that were above chance are marked with an orange asterisk mark
